# Stem and progenitor cell proliferation are independently regulated by cell type-specific *cyclinD* genes

**DOI:** 10.1101/2024.10.21.619490

**Authors:** Mark E. Lush, Ya-Yin Tsai, Shiyuan Chen, Daniela Münch, Julia Peloggia, Jeremy E. Sandler, Tatjana Piotrowski

## Abstract

Regeneration and homeostatic turnover of solid tissues depend on the proliferation of symmetrically dividing adult stem cells, which either remain stem cells or differentiate based on their niche position. Here we demonstrate that in zebrafish lateral line sensory organs, stem and progenitor cell proliferation are independently regulated by two *cyclinD* genes. Loss of *ccnd2a* impairs stem cell proliferation during development, while loss of *ccndx* disrupts hair cell progenitor proliferation but allows normal differentiation. Notably, *ccnd2a* can functionally replace *ccndx*, indicating that the respective effects of these Cyclins on proliferation are due to cell type-specific expression. However, even though hair cell progenitors differentiate normally in *ccndx* mutants, they are mispolarized due to *hes2* and Emx2 downregulation. Thus, regulated proliferation ensures that equal numbers of hair cells are polarized in opposite directions. Our study reveals cell type-specific roles for *cyclinD* genes in regulating the different populations of symmetrically dividing cells governing organ development and regeneration, with implications for regenerative medicine and disease.

## Introduction

Tissue turnover and regeneration are essential for organismal function and survival and require the proliferation of stem cells. In most solid tissues, this process relies on symmetrically dividing adult stem cells [1]. For instance, in the epithelia of the intestine, stomach, esophagus and the skin, stem cells divide symmetrically and—depending on their location in the stem cell niche—the daughter cells either maintain their stem cell characteristics, or if displaced from the niche proceed to differentiate [2–9]. Stem and progenitor cell proliferation need to be tightly controlled, as misregulation of niche signals or uncontrolled proliferation of stem and daughter cells can lead to serious diseases such as cancer. Despite the importance of symmetrically dividing stem cells and their progeny for tissue maintenance, regeneration and disease, whether their proliferation is differentially regulated has not been explored.

The zebrafish sensory lateral line is an excellent model to study the regulation of sensory organ homeostasis and regeneration at the cellular and molecular level within single cells [10–12]. It consists of clusters of 50-80 cells, called neuromasts that are arranged in lines on the head and along the trunk of the fish (Figure 1A-1C). These neuromasts are deposited during embryonic development by migrating primordia [13]. Each neuromast contains mechanosensory hair cells surrounded by support cells and peripheral mantle cells (Figures 1C-1D). Hair cells possess a long, microtubule based kinocilium and shorter actin rich stereocilia that are sensitive to water motion (Figure 1B, [14]).

**Figure 1.**
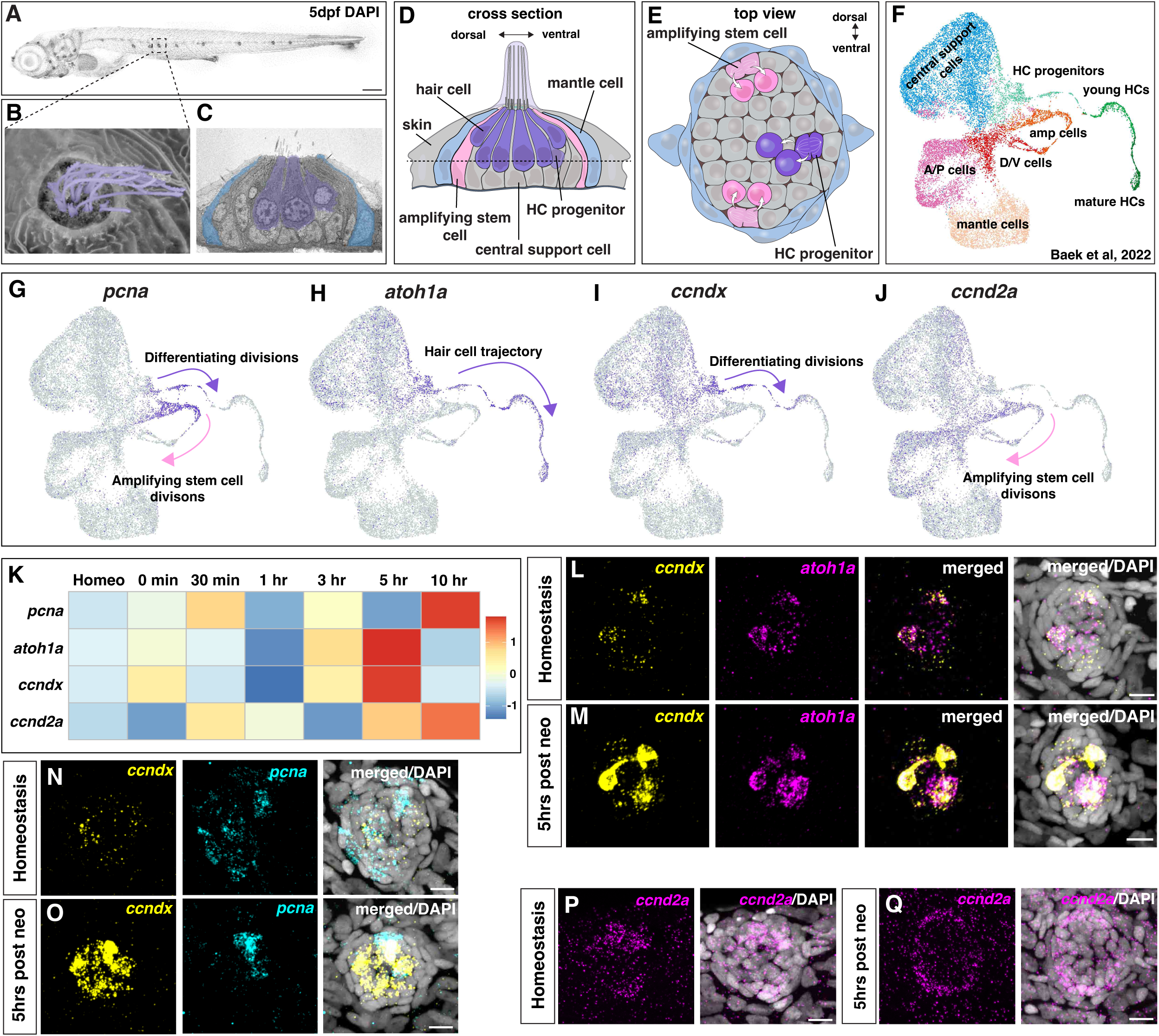
*ccndx* and *ccnd2a* are dynamically expressed in different proliferating cells in the regenerating zebrafish lateral line. (A) 5dpf DAPI-stained zebrafish larva, posterior lateral line neuromast in boxed region. Scale bar= 200μm. (B) Scanning electron micrograph of a 5dpf zebrafish neuromast (dorsal view) with short stereocilia and long kinocilia in purple (modified after Lush and Piotrowski, 2014 [10]). (C) Transmission electron micrograph of a transverse section of a 5dpf neuromast with hair cells in purple and mantle cells in blue. Additional support cells are unlabeled (modified after Lush and Piotrowski, 2014 [10]). (D and E) Diagram of a neuromast showing a transverse section (D) and a dorsal view (E). Progenitor cells and hair cells are in purple, amplifying stem cells in the dorsal-ventral poles in pink and mantle cells in blue. (F) Integrated scRNA-seq UMAP plot of a neuromast regeneration time course (homeostasis, 0min, 30min, 1h, 3h, 5h and 10h after hair cell death; Baek et al., 2022 [12]). (G-J) scRNA-seq Feature Plots (Baek et al., 2022 [12], https://piotrowskilab.shinyapps.io/neuromast_regeneration_scRNAseq_pub_2021/) illustrating gene-specific expression patterns. (G) *pcna* labels both dividing, differentiating hair cell progenitors (purple arrow) and amplifying stem cells (pink arrow). (H) *atoh1a* is expressed in some central cells and marks the lineage from hair cell progenitors to hair cells (purple arrow). (I) *ccndx* is expressed in some central cells and along the hair cell lineage and is highest in progenitor cells undergoing differentiating divisions (purple arrow). Its expression is quickly shut off as hair cells mature. (J) *ccnd2a* is more broadly expressed but is highest in the amplifying cell population (pink arrow) and absent from the hair cell lineage. (K) Heatmap of scaled gene expression during the regeneration time course (Baek et al., 2022 [12]), showing the same genes as in (G-J). All genes are briefly upregulated at 0-30 min but show the largest activation between 3-10 hrs. *atoh1a* and *ccndx* show similar expression dynamics, whereas *ccnd2a* expression peaks slightly later. (L and M) HCR in situ hybridization of *ccndx* (yellow) and *atoh1a* (magenta) during homeostasis (L) and 5 hrs after hair cell death (M). Scale bar= 10μm. (N and O) HCR of *ccndx* (yellow) and *pcna* (cyan) during homeostasis (N) and 5 hrs after hair cell death (O). *ccndx* and *pcna* show co-localization in a few cells and the expression of both is increased 5 hrs after neomycin treatment. Scale bar= 10μm. (P and Q) HCR of *ccnd2a* (magenta) at homeostasis (P) and 5 hrs post hair cell death (Q). *ccnd2a* expression becomes broader, extending into the anterior-posterior poles after hair cell death. Scale bar= 10μm.

Homeostasis and regeneration of neuromast cells are maintained by two populations of proliferating cells: amplifying support cells and differentiating hair cell progenitors (Figure 1E, [15]). In the dorsal-ventral (D-V) poles, support cells (Figure 1E, pink cells) divide and their daughter cells can adopt different fates: remain undifferentiated while in contact with mantle cells, the hypothetical niche, or, be displaced away from the niche and become a hair cell progenitor [15, 16]. Because amplifying cells self-renew and give rise to hair cells, we call them amplifying stem cells. Their degree of plasticity or potency to generate other neuromast cell types remains to be tested. Hair cell progenitors in the center of the organ will in turn divide again and give rise to two hair cells (Figure 1E, purple cells, [10, 15, 17, 18]). After hair cell loss progenitors also arise from non-D-V pole support cells, demonstrating plasticity, similar to other epithelial cell lineages [19, 20].

As in other mechanosensory organs in various species, zebrafish hair cell differentiation in neuromasts is negatively regulated via Notch-dependent lateral inhibition, and loss of Notch signaling causes the development of more hair cells at the expense of support cells [15, 17, 18, 21, 22]. In addition to its function in progenitor fate specification, Notch signaling also inhibits proliferation of differentiating progenitor cells during regeneration [15, 17]. Therefore, during regeneration and immediately after hair cell death, Notch signaling is downregulated, leading to differentiation of progenitor cells, their proliferation and further differentiation into hair cells. The current belief is that cell proliferation is essential for neuromast hair cell regeneration, as cell cycle inhibition with pharmacological inhibitors leads to a failure in regeneration [18, 23].

Moreover, progenitor cell division produces a pair of hair cells with opposing polarity, ensuring the equal generation of hair cells that are sensitive to either rostrad or caudad water flow [24]. The transcription factor Emx2 determines hair cell polarity in both the lateral line and ear [25–30]. In neuromasts, it is expressed in only one of the two sibling hair cells, where it reverses the cell’s default anterior polarity. Notch-mediated lateral inhibition between the two initially equal progenitors inhibits the expression of Emx2 in one progenitor, and loss of Notch signaling causes both hair cells in the pair to acquire the same polarity [25, 28, 30].

As proliferation is not only essential for life-long regeneration but also for correct hair cell polarity it is essential to elucidate how it is controlled. We previously characterized gene expression dynamics during regeneration in all neuromast cell types using single cell RNA-seq (scRNA-seq) [11, 12]. Here we show that the proliferating amplifying stem cell and progenitor cell populations of the zebrafish lateral line express different *cyclinD* genes, which are G1-regulators that bind and activate cyclin-dependent kinases (CDK4/6) to initiate the cell cycle [31, 32]. Amplifying cells express *ccnd2a* and dividing, differentiating progenitor cells express *ccndx.* Loss of *ccndx* causes lack of differentiating progenitor divisions while leaving amplifying divisions unaffected. Hair cells still regenerate in lower numbers, demonstrating that progenitor cell proliferation is not required for differentiation and regeneration of hair cells. In contrast, loss of *ccnd2a* only affects amplifying cell divisions, at least during development. *ccnd2a* driven by the *ccndx* promoter rescues progenitor proliferation in *ccndx* mutants, demonstrating that the cell type-specific effects of these two D-type cyclins are caused by their cell type-specific expression, not because they interact with different targets. Thus, proliferation in amplifying cells and differentiating progenitor cells is mechanistically uncoupled. We also show that Notch signaling inhibits *ccndx* during homeostasis and that the increase in hair cell progenitor proliferation after Notch downregulation during hair cell regeneration requires *ccndx*. Lastly, we demonstrate using scRNA-seq and functional analyses that loss of *ccndx* and progenitor proliferation lead to hair cell polarity defects due to *hes2* downregulation and ectopic Emx2 expression. Our findings have important implications for the understanding of how proliferation of symmetrically dividing stem and progenitor cells is controlled during homeostasis and disease.

## Results

### Zebrafish lateral line sensory organs possess two populations of dividing cells with unique gene expression profiles

To uncover the mechanisms underlying hair cell regeneration, we set out to identify new regulators of this process. In zebrafish, the antibiotic neomycin induces rapid hair cell death followed by complete regeneration [10, 33, 34]. We used this approach in our previous scRNA-seq time course, where we identified all cell types of the regenerating neuromasts (Figure 1F, [12]). With regard to dividing cells, two trajectories of proliferating populations were found to be marked by *pcna* expression (Figure 1G; [12]): the dividing hair cell progenitor cells mature into hair cells and are marked by the hair cell-specifying transcription factor *atoh1a*, whereas amplifying cells are defined as *pcna^+^* cells that do not express *atoh1a* or hair cell genes (Figure 1H, [12]).

To identify additional genes specific to either cell population, we queried the scRNA-seq data sets our lab previously published [11, 12]. We found the *cyclinD* family member *ccndx* to be specifically expressed in differentiating progenitor cells (differentiating divisions, Figure 1I), whereas *ccnd2a* was enriched in amplifying cells (amplifying divisions, Figure 1J). Both *cyclinD* genes, as well as *atoh1a* and *pcna* were differentially expressed during regeneration with *atoh1a* and *ccndx* possessing similar expression dynamics (Figure 1K and Figure S1A-S1H, [12, 15, 17, 33]. Upon neomycin treatment, all genes were initially downregulated followed by a strong upregulation between 3-10 hours (hrs), when cells started to proliferate (Figure 1K, [15]). We validated the scRNA-seq time course data with hybridization chain reaction (HCR) mRNA expression analyses. *ccndx* and *atoh1a* were co-expressed in progenitor cells during homeostasis and both genes showed strong upregulation at 5 hrs post hair cell death (Figures 1L-1M). *pcna* and *ccndx* overlapped in a few cells during homeostasis and 5 hrs after neomycin treatment (Figure 1N-1O). *ccndx* was also expressed in hair cell progenitor cells in the 32 hours post fertilization (hpf) migrating lateral line primordium (Figure S2A) and spinal cord neurons as described in the frog and zebrafish (Figure S2B [35, 36]), but was absent from the 5 days post fertilization (dpf) zebrafish ear (Figures S2C). In contrast to *ccndx*, *ccnd2a* was enriched in amplifying cells during homeostasis, with broader expression 5 hrs after hair cell death (Figures 1P-1Q and Figure S1D and S1H).

In conclusion, two *cyclinD* genes are expressed in amplifying stem cells and proliferating hair cell progenitor cells, respectively, suggesting that proliferation is differentially regulated in stem cells and their progeny.

### *ccndx*^-/-^ neuromasts form new hair cells through direct differentiation of progenitor cells in the absence of proliferation

To test the function of *ccndx*, we generated a mutant using CRISPR/Cas9 (Figure S3A). The mutation introduces a stop codon, but truncated *ccndx* mRNA is still transcribed in these mutants (Figures S3F and S3G). 5 dpf *ccndx^-/-^* larvae looked grossly normal but lacked an inflated swim bladder and showed abnormal swimming behavior (Figures S3B and S3C). Nevertheless, the migrating lateral line primordium deposited neuromasts normally during embryogenesis (Figures S3D and S3E). Previous morpholino-induced knockdown of *ccndx* in zebrafish and frogs resulted in defects in motor neuron formation and axonal outgrowth [35, 36], however anti-acetylated tubulin staining in our 5 dpf *ccndx^-/-^* larvae showed no obvious differences in motor axon outgrowth (Figures S3H-S3K). To test the role of *ccndx* in hair cell formation, we crossed *ccndx^-/-^* zebrafish with a hair cell-specific reporter line (*Tg(myo6b:H2B-mScarlet-I).* We observed a reduced number of hair cells in 5 dpf homeostatic *ccndx^-/-^* neuromasts and fewer regenerated hair cells 24- and 48 hrs after neomycin-induced hair cell death compared to sibling larvae (Figures 2A-2C).

**Figure 2.**
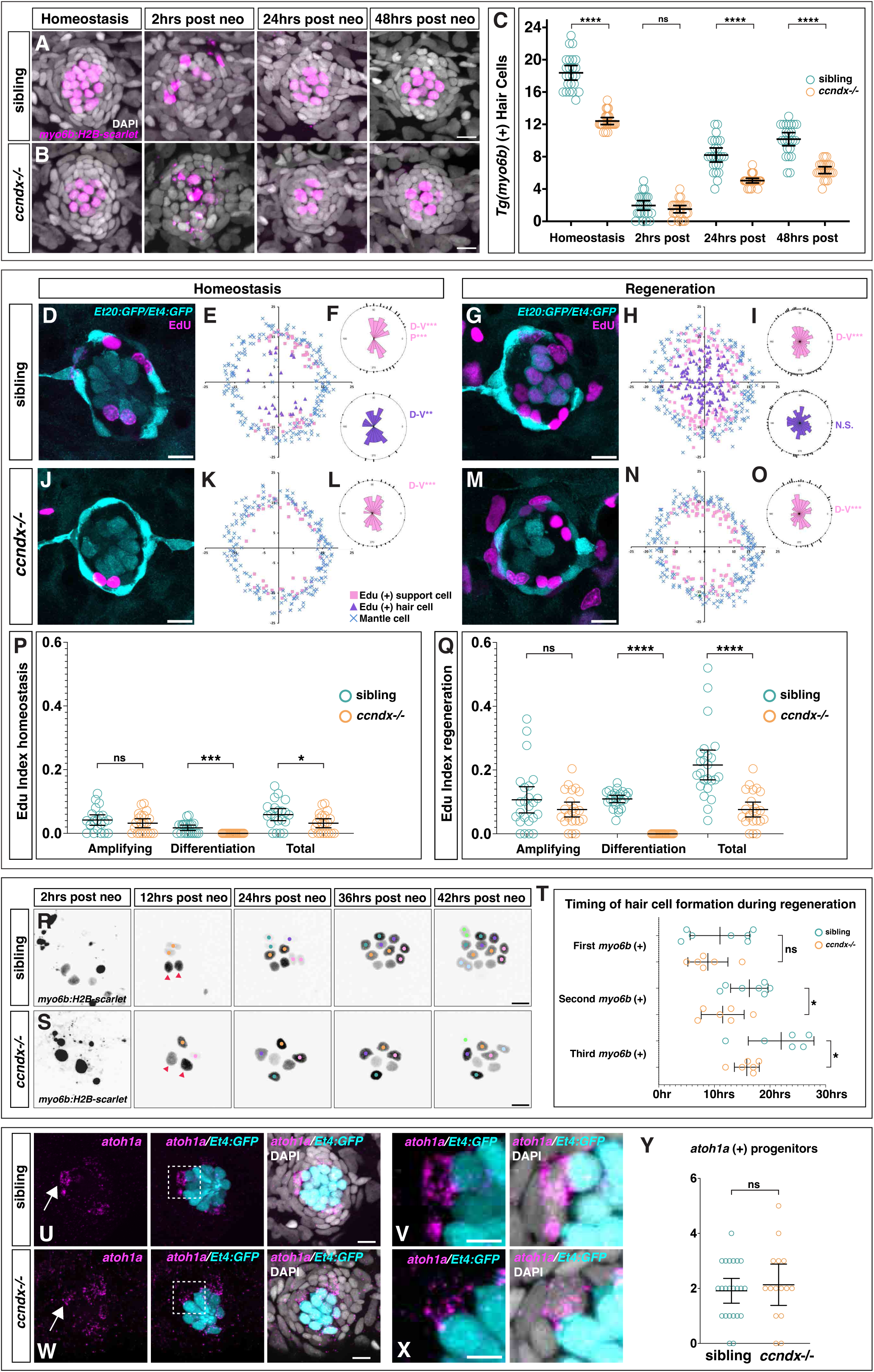
In *ccndx* mutants hair cells regenerate through direct differentiation in the absence of proliferation. (A and B) Time course of hair cell regeneration in sibling (A) and *ccndx^-/-^* (B) at homeostasis and 2, 24 or 48 hrs after hair cell death. Hair cells are labeled by expression of *myo6b:H2B-mScarlet-I* (magenta). Scale bar= 10μm. (C) *myo6b:H2B-mScarlet-I*^+^ hair cell counts in sibling and *ccndx^-/-^* at homeostasis and 2, 24 or 48 hrs after hair cell death. (D-F) (D) Representative image of a sibling neuromast expressing *sqEt4:GFP/sqEt20:GFP* stained with EdU (magenta) after 5-6 dpf of homeostasis. Scale bar= 10mm. (E) Spatial analysis of multiple neuromasts showing EdU^+^ support cells (pink squares), EdU^+^ hair cells (purple triangles) and surrounding mantle cells (blue X’s). (F) Statistical analysis of position of EdU^+^ support cells (pink) and hair cells (purple). (G-I) Representative image of a sibling neuromast 24 hrs after neomycin treatment expressing *sqEt4:GFP/sqEt20:GFP* and stained with EdU (magenta) (G). Scale bar=10μm. Spatial analysis of multiple sibling regenerating neuromasts (H) and statistical analysis of EdU^+^ cell positions (I). (J-L) Representative image of a *ccndx^-/-^* neuromast expressing *sqEt4:GFP/sqEt20:GFP* stained with EdU (magenta) after 5-6 dpf of homeostasis (J). Scale bar= 10 μm. Spatial analysis of multiple *ccndx^-/-^* homeostatic neuromasts (K) and statistical analysis of EdU^+^ cell positions (L). No EdU^+^ hair cells are present. (M-O) Representative image of a *ccndx^-/-^* neuromast 24 hrs after neomycin treatment expressing *sqEt4:GFP/sqEt20:GFP* and stained with EdU (magenta) (M). Scale bar= 10μm. Spatial analysis of multiple *ccndx^-/-^* regenerating neuromasts (N) and statistical analysis of EdU^+^ cell positions (O). No EdU^+^ hair cells are present. (P) EdU indexes of sibling and *ccndx^-/-^* amplifying, differentiation or total Edu+ cells during homeostasis. There is no difference in the amplifying index but there are no EdU^+^ hair cells in *ccndx^-/-^*. (Q) EdU indexes of sibling and *ccndx^-/-^* amplifying, differentiation and total EdU+ cells after 24 hrs of regeneration. The amplifying index is unaffected but *ccndx^-/-^* lack EdU^+^ hair cells. (R) Time-lapse analysis of regenerating sibling *myo6b:H2B-mScarlet-I*^+^ hair cells from 2 to 42 hrs after neomycin treatment. Red triangles point to hair cells that survived neomycin. Each new pair of hair cells is marked by differently colored dots. Scale bar= 10 μm. (S) Time-lapse analysis of regenerating *ccndx^-/-^ myo6b:H2B-mScarlet-I*^+^ hair cells from 2 to 42 hours after neomycin treatment. Red triangles point to hair cells that survived neomycin. Each new hair cell is marked by differently colored dots. (T) Quantification of the appearance of the first, second and third *myo6b:H2B-mScarlet-I*^+^ hair cell in sibling and *ccndx^-/-^* neuromasts over 30 hrs of regeneration. For siblings, we counted the appearance of each new *myo6b:H2B-mScarlet*^+^ cell before division. (U and V) (U) *atoh1a* HCR (magenta) and *sqEt4:GFP*^+^ (cyan) hair cells in 5dpf sibling neuromasts. (V) A magnified *atoh1a*^+^/*sqET4:GFP*^-^ progenitor cell from (U). Scale bar= 10 μm. (W and X) (W) *atoh1a* HCR (magenta) and *sqEt4:GFP*^+^ (cyan) hair cells in 5dpf *ccndx^-/-^* neuromasts. (X) A magnified *atoh1a*^+^/*sqET4:GFP*^-^ progenitor cell from (W). Scale bar= 10 μm. (Y) Quantification of 5 dpf *atoh1a*^+^/*sqET4:GFP*^-^ progenitor cells showing no difference between sibling and *ccndx^-/-^*.

As hair cell regeneration in the zebrafish lateral line is thought to depend on progenitor proliferation [18, 23], we examined whether proliferation was affected in *ccndx^-/-^* neuromasts during homeostasis and regeneration by assessing EdU incorporation for 24 hrs, which labels cells that have entered S-phase of the cell cycle. To determine if the location of amplifying and differentiating divisions was affected by loss of *ccndx*, we performed spatial analyses of sibling and mutant neuromasts by plotting the location of all EdU^+^ support and hair cells from multiple neuromasts [15, 37]. EdU+ cells that also express *sqEt4:GFP* represent differentiating hair cells (purple triangles; Figures 2E, 2H, 2K and 2N), while EdU^+^ cells that are not *sqEt4GFP^+^* are amplifying cells, plotted as pink squares. Quiescent *sqEt20^+^* mantle cells are plotted as blue crosses and delineate the neuromast outline [15, 37].

In homeostatic sibling larvae, amplifying cell divisions were restricted to the D-V poles of neuromasts, while differentiating cell divisions were centrally located (Figure 1E and Figure 2D-2F, [15]). Upon regeneration, the rate of cell division significantly increased in both cell populations (Figure 2G-2I). In homeostatic and regenerating *ccndx^-/-^* larvae, amplifying divisions were also restricted to the D-V poles and EdU indexes were not significantly different from siblings (Figures 2J-2Q). In stark contrast, there were no EdU^+^ hair cells present in *ccndx^-/-^* neuromasts (Figures 2J-2O). Quantification of cell divisions showed that the only dividing cells in *ccndx*^-/-^ neuromasts were the D-V amplifying cells (Figures 2P and 2Q). The decrease in total EdU index in *ccndx^-/-^* neuromasts is therefore solely due to the lack of EdU^+^ hair cells.

To visualize hair cell formation *in vivo,* we performed time-lapse analyses of regenerating neuromasts expressing *myo6b:H2B-mScarlet-I. myo6b:H2B-mScarlet-I* started to be expressed early in hair cell differentiation, just before the progenitor divided (Supplemental Movie 1, Figure 2R). In contrast, *ccndx^-/-^* mutant progenitor cells started to express *myo6b:H2B-mScarlet* but failed to divide (Supplemental Movie 1, Figure 2S). Thus, only a single hair cell was generated per progenitor cell compared to two hair cells in siblings. These results challenge the current paradigm, showing that neuromasts can regenerate hair cells without progenitor proliferation.

The cell cycle affects differentiation in several cell types [38–41] and we therefore asked if the timing of hair cell differentiation during regeneration differs in the absence of proliferation in *ccndx*^-/-^ neuromasts. We analyzed time-lapse recordings of the first 30h of regeneration of *myo6b:H2B-mScarlet-I^+^* hair cells in sibling and *ccndx* mutant neuromasts. The appearance of the first new hair cells was not significantly different between siblings and *ccndx* mutant animals, whereas the subsequent second and third hair cells appeared significantly faster in *ccndx^-/-^* larvae (Figure 2T, Supplemental Movie 1). RNA transcription is greatly decreased as cells go through mitosis [42], hence it may be possible that expression of differentiation genes proceeds faster in cells that do not go through the cell cycle.

As shown in the time lapse movies and EdU experiments, *ccndx* is required for progenitor proliferation, and we wondered if loss of *ccndx* might affect how many hair cell progenitors are generated even before they divide. To examine progenitor cell numbers during homeostasis we performed HCRs for *atoh1a* in neuromasts expressing *sqEt4:GFP* at 5dpf. We considered progenitor cells as those without *sqEt4:GFP* ^-^ but strongly expressing *atoh1a*^+^. The number of *atoh1a*^+^/*sqEt4:GFP* ^-^ cells in *ccndx^-/-^* neuromasts was not significantly different from siblings (Figures 2U-2X), illustrating that progenitor numbers were normal during homeostasis in *ccndx^-/-^* larvae (Figure 2Y). The analysis of hair cell regeneration in *ccndx*^-/-^ larvae demonstrates that hair cells can regenerate via direct differentiation of progenitor cells in the absence of proliferation. There are fewer *ccndx^-/-^* hair cells compared to siblings because mutant progenitor cells generate one hair cell instead of two during homeostasis.

The finding that *ccndx^-/-^* hair cells differentiate in the absence proliferation contrasts with previous reports that showed pharmacological cell cycle blockers inhibit neuromast hair cell regeneration [18, 23]. To investigate the reason for this discrepancy, we treated 5 dpf control and *ccndx^-/-^* larvae with the early S-phase inhibitor aphidicolin and EdU during 48 hrs of regeneration. As published, control larvae treated with aphidicolin showed a severe reduction in regenerated hair cells compared to DMSO controls (Figure S4A, S4B and S4E). Aphidicolin-treated *ccndx^-/-^* regenerated more hair cells than treated siblings (Figure S4C, S4D and S4E). In aphidicolin-treated siblings, EdU indexes showed a clear reduction in the amplifying and differentiating populations, and therefore total proliferation (Figures S4F-S4H). EdU^+^ support cells were also reduced to the same extent by aphidicolin in *ccndx^-/-^*, but as *ccndx^-/-^* normally do not produce EdU^+^ hair cells, aphidicolin had no added effect on this population (Figures S4F-S4H). Aphidicolin-treated siblings often showed enlarged hair cells, as if they were trying to divide but then failed (Figure S4B, yellow asterisk). This result was also seen in time-lapse analyses (Supplemental Movie 2). In sibling neuromasts, hair cells began to differentiate as evidenced by upregulation of *myo6b:H2B-mScarlet-I*. However, the newly generated hair cells failed to divide, instead enlarged and died. In contrast, aphidicolin-treated *ccndx^-/-^* hair cells differentiated and survived over the 42 hrs of imaging. We conclude that in aphidicolin-treated siblings, new hair cells start to differentiate but eventually die, possibly via activation of cell cycle check-point regulators. As *ccndx^-/-^* progenitor cells fail to initiate the cell cycle they escape the toxicity of aphidicolin treatment. Our experiments hence show that proliferation is not required for zebrafish lateral line hair cell regeneration and that previous studies using aphidicolin to stop the cell cycle caused regeneration defects due to drug-induced hair cell death.

### *ccnd2a* drives amplifying cell divisions but substitutes for *ccndx* if expressed in progenitor cells

In the amplifying stem cell population, *ccnd2a* is the most enriched *cyclinD* gene and proliferation of this cell population is not affected in *ccndx^-/-^* mutants (Figures 1J and 1K). To test the function of *ccnd2a* we generated a CRISPR-Cas12a deletion mutant (Figure S5A)*. ccnd2a^-/-^* larvae are grossly normal, survive to adulthood and are fertile (Figures S5B and S5C). However, neuromasts fail to grow between 48-72 hpf after they are deposited by the migrating primordium (Figures 3A-3G, Figures S5D and S5E). To test if the smaller neuromasts are caused by a decrease in proliferation during development, we treated *ccnd2a^+/+^* and *ccnd2a^-/-^* with EdU for 24 hrs from 2-3 dpf. In *ccnd2a*^-/-^ neuromasts amplifying divisions were significantly reduced, consistent with *ccnd2a* expression being enriched in these cells, whereas hair cell differentiation divisions were only slightly increased (Figure 3H).

**Figure 3.**
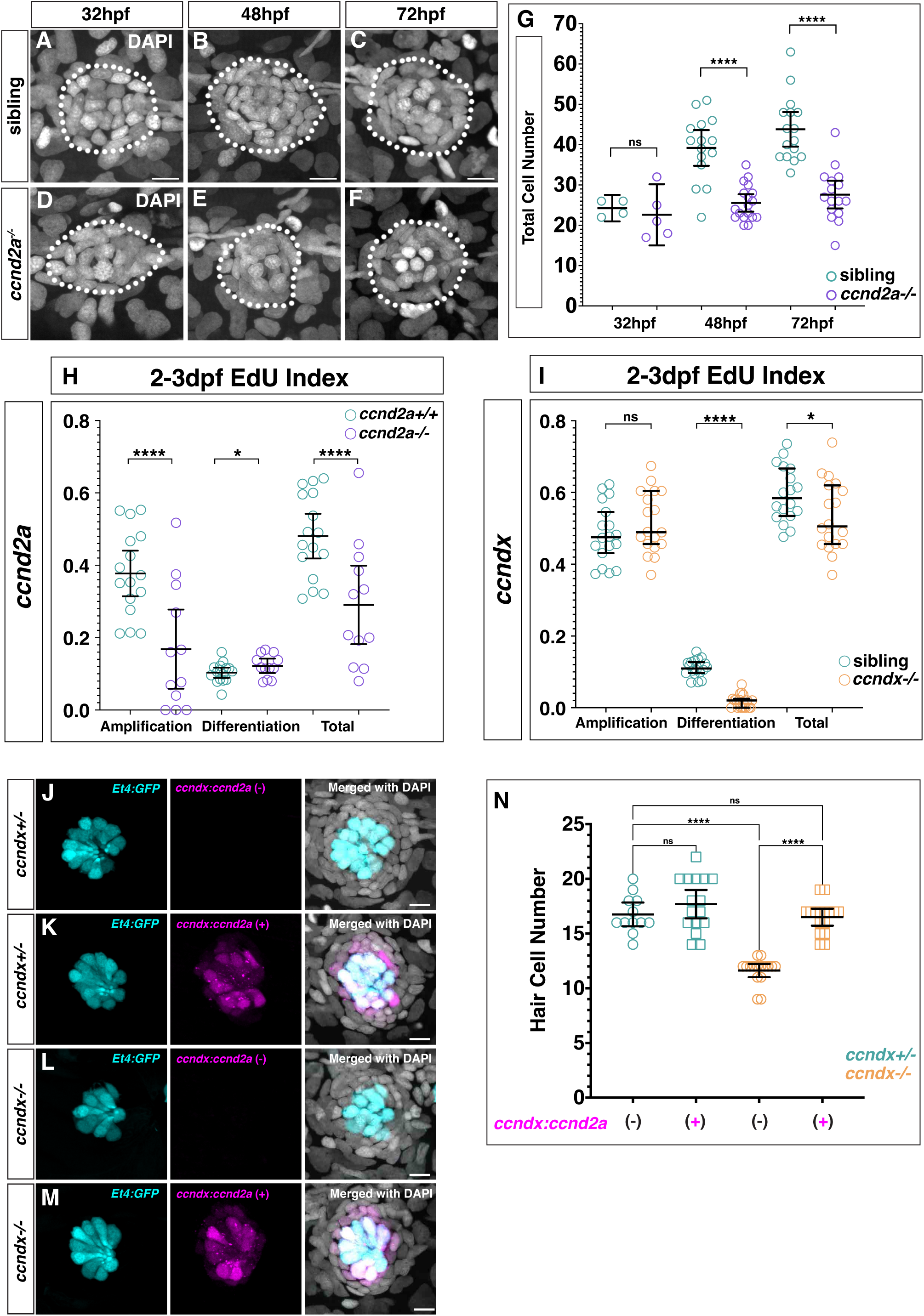
*ccnd2a* and *ccndx* are required for support cell and progenitor cell proliferation during development, respectively. (A-F) DAPI stained nuclei of neuromasts from sibling (A-C) or *ccnd2a^-/-^* (D-F) at 32, 48 or 72 hpf. Scale bar= 10 μm. (G) Quantification of total cell number from DAPI stained neuromasts in sibling and *ccnd2a^-/-^* at 32, 48 or 72 hpf. (H) Comparison of wildtype and *ccnd2a^-/-^* amplifying, differentiation or total EdU indexes between 2-3 dpf. *ccnd2a^-/-^* has reduced amplification and total EdU indexes and a slightly increased differentiation index. (I) Comparison of sibling and *ccndx^-/-^* amplifying, differentiation or total EdU indexes between 2-3 dpf. Similarly to 5-6 dpf, *ccndx^-/-^* neuromasts show reduced differentiation and total EdU indexes and no change in the amplification index. (J-M) Representative images of 5dpf neuromasts from *ccndx^+/-^* (J and K) or *ccndx^-/-^* (L and M) expressing *sqEt4:GFP* without or with *myo6b:H2B-mScarlet-I* driven by the *ccndx* promoter. Scale bar= 10 μm. (N) Quantification of *sqEt4:GFP*^+^ hair cells in *ccndx^+/-^* or *ccndx^-/-^* neuromasts without or with *ccnd2a-P2A-mScarlet-I*.

We found that *ccnd2a* is only required for amplifying divisions during development but not regeneration, as after injury, amplifying proliferation still occured in the D-V poles of *ccnd2a^-/-^*(Figures S5F-S5K). This finding reveals that amplifying stem cell proliferation is differentially regulated during development and regeneration, and opens the possibility that another *cyclinD,* such as *ccnd1* may functionally compensate during regeneration.

In contrast to *ccnd2a^-/-^*, in 2-3 dpf developing *ccndx^-/-^* neuromasts amplifying proliferation is unaffected, and only differentiating divisions are decreased (Figure 3I). Thus, *ccndx* is required for hair cell progenitor proliferation during development, homeostasis and regeneration, whereas *ccnd2a* is necessary for amplifying stem cell divisions only during development.

As *ccndx* and *ccnd2a* control proliferation of distinct proliferating cell populations we wondered if they possess different, cell cycle-independent functions in the two cell types. To test the specificity of these genes we expressed a *ccnd2a-P2A-mScarlet-I* transgene under the control of *ccndx* regulatory sequences (described below) in *ccndx*^-/-^ and sibling larvae. *ccnd2a* expression does not increase hair cell numbers in *ccndx^+/-^* but rescues hair cell numbers in *ccndx^-/-^* neuromasts (Figure 3J-3N). This demonstrates that *ccndx* does not possess specific functions in the regulation of hair cell formation. The cell type-specific effects of *ccndx* are therefore a result of its spatially and temporally restricted expression pattern only in the progenitor cell population.

### *atoh1a* is dispensable for *ccndx* expression in hair cell progenitors

To better define the sequence of signaling events leading to proliferation of hair cell progenitors we aimed to identify possible upstream regulators of *ccndx*. To identify regulatory sequences that drive *ccndx* expression, we compared zebrafish and goldfish genomes [43, 44], which revealed two conserved regions upstream of *ccndx* (Figure 4A). We generated reporter lines driving EGFP or destabilized EGFP via these elements, cloning a 4.5kb fragment encompassing both conserved regions. By assaying reporter activity, we found that this 4.5kb region upstream of *ccndx* is sufficient to drive H2B-GFP expression in the hindbrain, spinal cord, migrating lateral line primordium and 5 dpf neuromasts (Figure 4B-4E).

**Figure 4.**
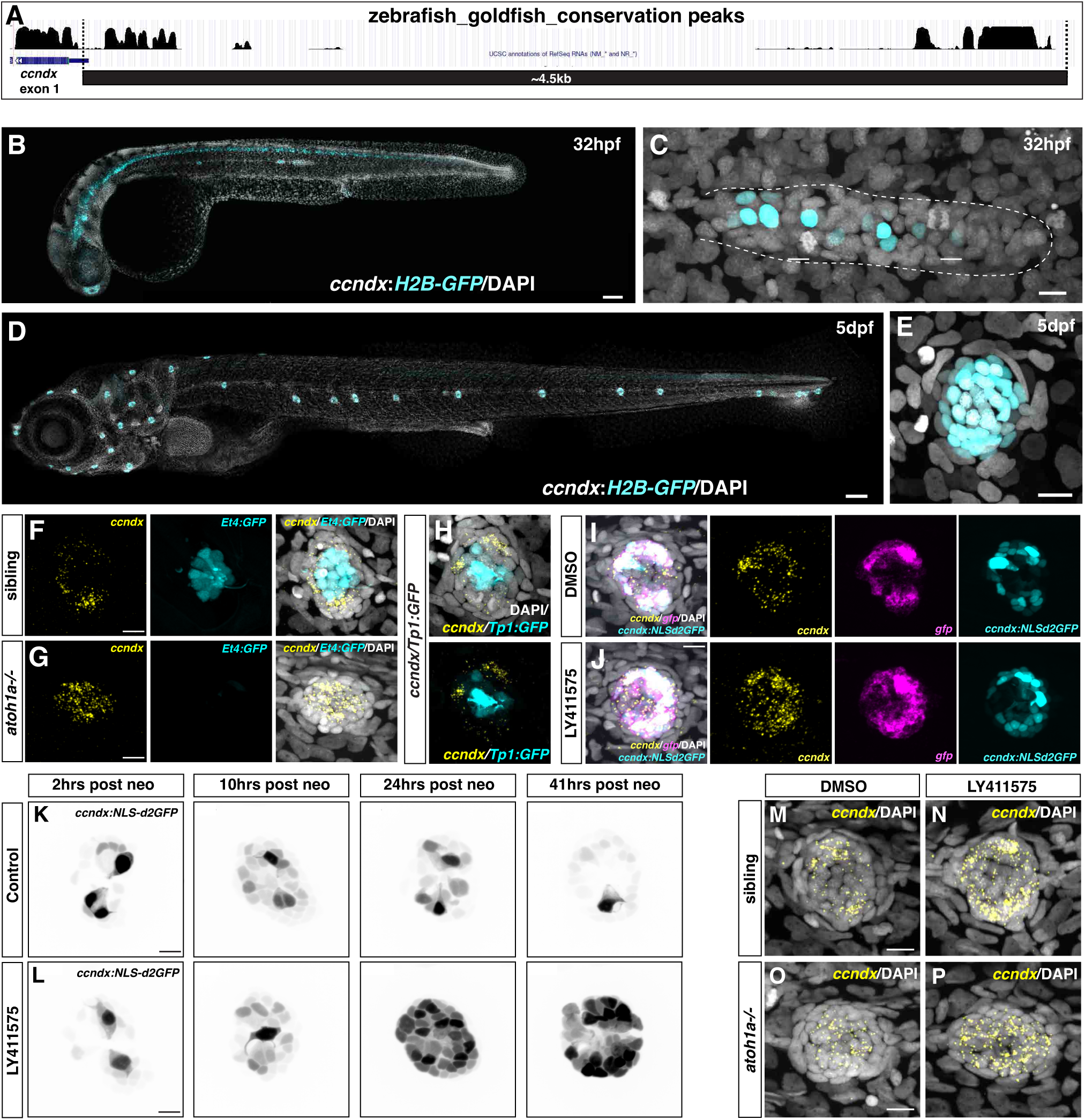
Characterization of *ccndx* enhancer regions and negative regulation of *ccndx* expression by Notch signaling. (A) UCSC zebrafish genome browser track showing two regions of conservation with the goldfish genome upstream of *ccndx* exon 1, showing the 4.5kb region cloned. (B and C) DAPI stained 32hpf zebrafish larvae showing H2B-EGFP (cyan) expression in the hindbrain, spinal cord and migrating lateral line primordium driven by the *ccndx* upstream region with a higher magnification image of the primordium (C). Scale bars= 10μm and 200 μm, respectively. (D and E) DAPI stained 5dpf zebrafish showing H2B-EGFP (cyan) expression in the spinal cord and neuromasts driven by the *ccndx* upstream region with higher magnification view of a neuromast (E). Scale bars=10μm and 200 μm, respectively. (F and G) *ccndx* HCR (yellow) in 5dpf sibling (F) or *atoh1a^-/-^* neuromasts (G). *sqEt4:EGFP* expression (cyan) shows lack of hair cells in *atoh1a^-/-^* larvae. *ccndx* is still expressed in *atoh1a^-/-^* neuromasts and shows a broader and more central expression pattern. Scale bar= 10μm. (H) *ccndx* HCR (yellow) in zebrafish expressing the *tp1bglobin:egfp* Notch reporter (cyan) showing the lack of co-localization. Scale bar= 10μm. (I and J) HCR for *ccndx* (yellow) and *egfp* (magenta) in *ccndx:NLSd2GFP* (cyan) transgenic zebrafish treated with DMSO (I) or LY411575 (J) for 6 hrs. LY411575 treatment induces an increase in *ccndx* and *egfp* expression, especially in the central region. *NLSd2GFP* fluorescence intensity is not changed at this timepoint. Scale bar= 10μm. (K and L) Images from time lapses of *ccndx:NLSd2EGFP* transgenic zebrafish which were treated with DMSO (K) or with LY411575 (L) immediately after neomycin treatment. There is an increase in the number of NLSd2EGFP^+^ cells around 10 hrs after neomycin which decreases in DMSO treated fish but continues to increase with Notch inhibition. Scale bar= 10μm. (M-P) HCR for *ccndx* (yellow) in 5dpf neuromasts in DMSO treated sibling (M) or *atoh1a-/-* (O) or LY411575 treated sibling (N) or *atoh1a^-/-^* (P). Scale bar= 10μm.

One possible regulator of *ccndx* is *atoh1a,* as it plays a crucial role in hair cell specification, is co-expressed with *ccndx* in progenitor cells and exhibits highly similar expression dynamics (Figures 1K-1M). An interaction between the two genes would imply a coupling of hair cell specification and proliferation. We therefore asked if *atoh1a* impacts *ccndx* expression by examining *atoh1a* mutant neuromasts. *ccndx* expression becomes broader in *atoh1a^-/-^* compared to siblings, showing a more central expression pattern (Figure 4F-4G). Therefore, *atoh1a* is not required for *ccndx* expression, and hair cell specification and *ccndx* expression are uncoupled in the lateral line.

### Notch signaling represses *ccndx* expression in hair cell progenitors

As Notch signaling inhibits hair cell progenitor proliferation during hair cell regeneration [15, 17], we examined whether Notch signaling inhibits *ccndx* expression. HCR of the Notch reporter line (*tp1bglobin:EGFP*) showed that *ccndx* and *egfp* are expressed in different cells (Figure 4H), suggesting that Notch signaling may indeed inhibit *ccndx* expression. To test if Notch represses *ccndx* we inhibited Notch signaling with the gamma-secretase inhibitor LY411575 in larvae expressing nuclear localized destabilized GFP driven by the *ccndx* promoter (*ccndx:NLSd2EGFP)*. After 6 hrs of Notch inhibition during homeostasis, both endogenous *ccndx* and *egfp* mRNA of the reporter were upregulated compared to untreated neuromasts, while the *NLSd2GFP* protein was unchanged (Figures 4I-4J). In time-lapse recordings of regenerating *ccndx:NLS-d2EGFP*/*myo6b:H2B-mScarlet-I* neuromasts we observed a strong increase in GFP fluorescence after Notch inhibition compared to control larvae (Figures 4K-4L and Supplemental Movie 3). This shows that the 4.5kb regulatory region of *ccndx* also responds to Notch inhibition during regeneration. In both DMSO and LY411575 treated *ccndx:NLS-d2GFP* neuromasts, GFP fluorescence intensity was increased 10 hrs after neomycin, likely in response to the downregulation of Notch signaling that occurs after hair cell death [12, 15, 17]. In control larvae, GFP expression then gradually decreased (Figure 4K and Supplemental Movie 4), whereas in LY411575 treated larvae *ccndx:NLS-d2GFP* fluorescence intensity continued to increase and stayed strong in support cells to the end of the time-lapse (Figure 4L and Supplemental Movie 4).

As inhibiting Notch signaling is known to increase *atoh1a* expression [15, 17], we wondered whether the increased *ccndx* expression after Notch inhibition might be due to increased *atoh1a* levels. We treated *atoh1a*^-/-^ zebrafish with LY411575 for 6 hrs and performed HCR for *ccndx*. The central *ccndx* expression in *atoh1a^-/-^* larvae is still increased after LY411575 treatment (Figure 4M-4P) demonstrating that *atoh1a* is not required for the upregulation of *ccndx* after Notch inhibition. Together, our results show that Notch signaling inhibits *ccndx* expression in neuromasts.

### Notch inhibition leads to *ccndx*-dependent progenitor proliferation

Downregulation of Notch signaling during hair cell regeneration increases progenitor proliferation but also hair cell specification at the expense of support cells due to its role in lateral inhibition [15, 17, 18, 25, 30]. To test if Notch signaling inhibits progenitor proliferation or hair cell specification via its repression of *ccndx,* we treated sibling and *ccndx^-/-^* larvae with LY411575 and EdU for 24 hrs following hair cell death. Spatial analyses of EdU^+^ cells in sibling neuromasts showed a significant increase in EdU^+^ hair cells after Notch inhibition as previously described (Figure 5A and 5C and 5I, [15]). The increase in differentiating divisions resulted in an increase in hair cells (Figure 5J). In *ccndx^-/-^* neuromasts treated with DMSO, no EdU^+^ hair cells are observed, and after Notch inhibition only very few EdU^+^ hair cells form (Figure 5E and 5G). This result is verified by a time-lapse analysis of a LY411575 treated *ccndx^-/-^* neuromast which shows no proliferation of hair cell progenitors (Supplemental Movie 5). Nevertheless, Notch inhibition results in the regeneration of more *sqEt4:GFP+* (but EdU^-^) hair cells in *ccndx^-/-^* neuromasts (Figure 5J). The increase in hair cell formation following Notch inhibition is accompanied by a decrease in support cell numbers in both siblings and *ccndx^-/-^* neuromasts (Figure 5K). In contrast to the differentiating divisions, the amplifying divisions occur normally in the D-V poles in the absence of Notch signaling in both siblings and mutants (Figures 5A-5D and 5E-5H and 5L). Together, these results demonstrate that in the absence of *ccndx,* Notch inhibition still promotes hair cell fate at the expense of support cell fate, and that therefore lateral inhibition occurs independently of *ccndx*. However, the increased progenitor proliferation induced by Notch inhibition in wildtype neuromasts is strictly dependent on *ccndx*.

**Figure 5.**
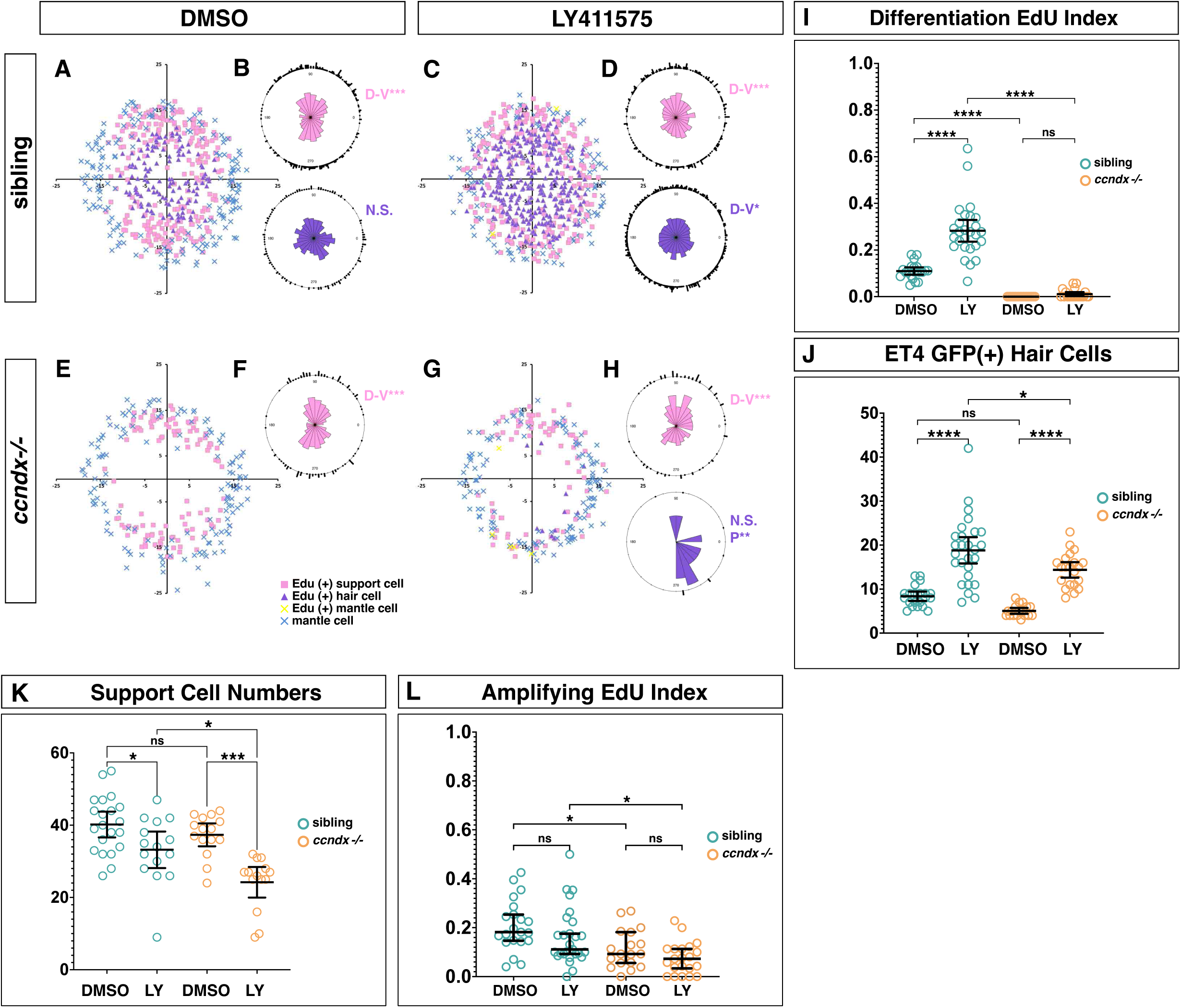
*ccndx* is required for the increased proliferation induced by Notch inhibition. (A and B) (A) Spatial analysis of EdU^+^ support cells and hair cells after 24 hrs of regeneration in control DMSO treated sibling neuromasts. (B) Statistical analysis of EdU^+^ cell positions. (C and D) (C) Spatial analysis of EdU^+^ support cells and hair cells after 24 hrs of regeneration in LY411575 treated sibling neuromasts. (D) Statistical analysis of EdU^+^ cell positions. (E and F) (E) Spatial analysis of EdU^+^ support cells and hair cells after 24 hrs of regeneration in control DMSO treated *ccndx^-/-^* neuromasts. (F) Statistical analysis of EdU^+^ cell positions. There are no EdU^+^ hair cells in *ccndx^-/-^* neuromasts. (G and H) (G) Spatial analysis of EdU^+^ support cells and hair cells after 24 hrs of regeneration in LY411475 treated *ccndx^-/-^* neuromasts. (H) Statistical analysis of EdU^+^ cell positions. There are few EdU^+^ hair cells in LY411575 treated *ccndx^-/-^* neuromasts. (I) EdU index of differentiation divisions after DMSO and LY411575 (LY) treatment in sibling and *ccndx^-/-^* regenerating neuromasts. There is no significant difference in differentiating EdU indexes between DMSO or LY411575 treated *ccndx^-/-^* neuromasts. (J) Quantification of *sqEt4:EGFP^+^* hair cells in DMSO and LY411575 (LY) treated sibling or *ccndx^-/-^* neuromasts after 24 hrs of regeneration. LY411575 induces an increase in regenerated hair cells in both siblings and *ccndx^-/-^* neuromasts. (K) Quantification of support cells in DMSO and LY411575 (LY) treated sibling or *ccndx^-/-^* neuromasts after 24 hrs of regeneration. LY411575 decreases support cells in both siblings and *ccndx^-/-^* neuromasts. (L) EdU index of amplifying cells in DMSO and LY411575 (LY) treated sibling or *ccndx^-/-^* neuromasts after 24 hrs of regeneration.

### *ccndx^-/-^* progenitors and hair cells differentiate normally but exhibit polarity defects

Hair cell progenitor divisions produce two hair cells of opposing polarity, resulting in neuromasts with 50% of hair cells possessing a kinocilium on one side and 50% with the kinocilium localized to the opposite side [24, 45]. To examine if hair cell polarity was affected in the absence of progenitor cell divisions we labeled hair cells with the F-actin binding peptide phalloidin. Siblings showed 50% of hair cells of either polarity, while in *ccndx^-/-^* neuromasts 70% of hair cells were polarized toward the posterior direction (Figure 6A-6C). This posterior bias in hair cell polarity in *ccndx^-/-^* neuromasts was rescued by *ccndx*-driven *ccnd2a* expression (Figure S6A-S6E), showing that the effect on hair cell polarity requires a *cyclinD* gene but is not specific to *ccndx*.

**Figure 6.**
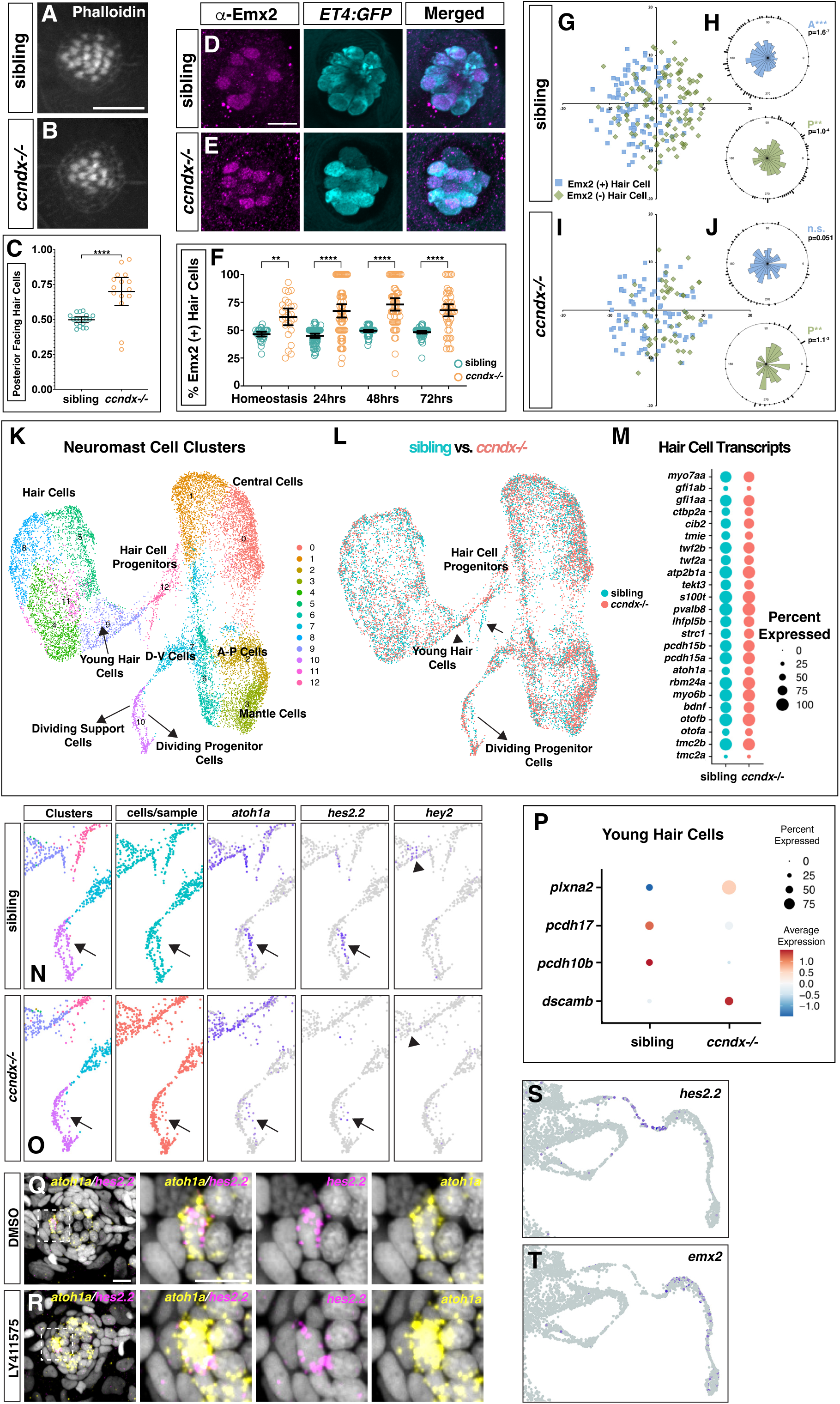
scRNA-seq of *ccndx^-/-^* neuromasts. (A-C) Phalloidin staining of sibling (A) and *ccndx^-/-^* (B) 5 dpf neuromasts with quantification of posterior-polarized hair cells per neuromast (C). Scale bar= 10μm. (D-F) Anti-Emx2 immunostaining (magenta) in 5dpf *sqEt4:EGFP* (cyan) expressing sibling (D) and *ccndx^-/-^* neuromasts (E). Scale bar=10μm. (F) Quantification of the percentage of anti-Emx2^+^ hair cells per neuromast during homeostasis and at 24, 48 or 72 hrs after neomycin treatment. (G-J) Spatial analysis of Emx2^+^ or Emx2^-^ hair cell nuclei cell position in sibling (G-H) and *ccndx^-/-^* (I-J) 72 hrs post neomycin treatment. (K) UMAP plot of integrated scRNA-seq datasets of 5dpf sibling and *ccndx^-/-^* lateral line cells with neuromast cell types labeled. (L) Same UMAP plot as in (K) with sibling cells (cyan) or *ccndx^-/-^* cells (red) individually labeled showing a reduction in the number of dividing progenitor cells (arrows) in *ccndx^-/-^* larvae. (M) Dot plot showing percentage of hair cells that express a given known hair cell gene comparing sibling (cyan) and *ccndx^-/-^* (red) mature hair cell populations. (N and O) Magnifications of scRNA-seq UMAP and Feature Plots of differentiating progenitor cells and young hair cells. Individual Feature Plots for *atoh1a*, *hes2.2*, and *hey2* in sibling (N) and *ccndx^-/-^* (O). *atoh1a* is expressed in progenitor cells and young hair cells. *hes2.2* is expressed in a subset of progenitor cells and is greatly reduced in *ccndx^-/-^* cells. *hey2* expression in young hair cells is reduced in *ccndx^-/-^*. (P) Dot plot of mRNA expression levels of a subset of differentially expressed genes comparing sibling and *ccndx^-/-^* young hair cell populations. (Q and R) HCR for *atoh1a* (yellow) and *hes2.2* (magenta) in a 5 dpf neuromast after DMSO (Q) or LY411575 (R) treatment showing co-expression within an individual progenitor cell. With cropped views of a single double positive cell. Scale bar=10μm. (S and T) Cropped image of *hes2.2* Feature Plot from scRNA-Seq regenerating time-course from Baek et al.,2022 [12] in progenitor cells (S) before *emx2* expression (T) in young hair cells.

Posteriorly polarized hair cells express the transcription factor Emx2 which is sufficient to drive posterior polarization in all hair cells if overexpressed [25, 28, 30, 46]. Antibody staining for Emx2 during homeostasis and at multiple time points after hair cell death showed that 70% of *ccndx^-/-^* hair cells express Emx2 compared to 50% in siblings, matching the phalloidin results (Figure 6D-6F). This higher percentage of Emx2^+^ hair cells persisted in *ccndx^-/-^* neuromasts as more hair cells were added during regeneration (Figure 6F). Spatial analyses of Emx2^+^ and Emx2^-^ hair cell nuclei revealed that hair cells are still spatially enriched in their respective halves of *ccndx^-/-^* neuromasts rather than being randomly distributed (Figures 6G-6J), showing that *ccndx^-/-^* hair cells continue to sense and react to the signals that polarize the field of hair cells across the sensory organ.

To determine if the changes in hair cell polarity are reflected by gene expression changes, we performed scRNA-Seq analyses of FAC-sorted neuromast cells from 5 dpf sibling and *ccndx^-/-^* larvae (Figures 6K and 6L, Table A). Sibling and *ccndx^-/-^* hair cells cluster together suggesting that their transcriptomes are similar. We examined the expression levels of candidate hair cell genes, including mechanotransduction related genes, and did not observe major differences between sibling and *ccndx^-/-^* hair cells (Figure 6M, Table B). We conclude that hair cell differentiation is not affected at a detectable manner by the absence of progenitor proliferation.

In contrast to mature hair cells, we observed more pronounced gene expression changes between sibling and *ccndx*^-/-^ hair cell progenitors and young hair cells (Figures 6L and 6N-6O, arrowheads). In siblings, *atoh1a*-expressing, dividing progenitor cells cluster closer with the amplifying cells, likely due to their expression of cell cycle genes (Figure 6L and 6N). In contrast, *ccndx^-/-^* hair cell progenitors form a continuous trajectory towards young hair cells (Figure 6L and 6O). Examining differentially expressed genes between siblings and *ccndx^-/-^ larvae* within these cell clusters which could be involved in regulating hair cell polarity, we found members of the *protocadherin*, *dscam* and *plexin* gene families (Figure 6P). These genes are expressed in either progenitor cells or young hair cells and could be involved in regulating hair cell polarity or hair cell rearrangements that coincide with the establishment of hair cell polarity [18, 47].

As Emx2 is inhibited by Notch signaling, we searched for gene expression differences in potential Notch pathway genes, as well as genes that are expressed immediately prior to *emx2* in progenitor cells. In *ccndx^-/-^* neuromasts, the Notch pathway members *her4.1*, *her15.2* and *hey2* are downregulated, as are *hes2.1, hes2.2*, which are expressed before *emx2* levels increase (Table B). As an example, in *ccndx^-/-^*, *hes2.2* is downregulated in hair cell progenitors (Figures 6N and 6O, arrows) while *hey2* is downregulated in young hair cells (Figures 6N and 6O, arrowheads). The downregulation of these genes suggests they could be involved in negatively regulating Emx2. HCR for *hes2.2* and *atoh1a* shows that *hes2.2* is expressed in some *atoh1a*+ cells (Figure 6Q), which is supported by the scRNA-seq regeneration time course that shows that *hes2.2* is transiently expressed in hair cell progenitor cells before they turn on *emx2* (Figures 6S and 6T, [12]). As *hes2.2* is not regulated by Notch signaling in neuromasts because LY411575 does not affect *hes2.2* expression (Figure 6R), the mechanism by which *hes2.2* expression is regulated by *ccndx* is Notch-independent.

### *ccndx* regulates hair cell polarity via activating *hes2.2* which inhibits *emx2*

To test if the loss of *hes2.2 expression* in *ccndx^-/-^* progenitor cells causes hair cell polarity defects, we deleted it with CRISPR-Cas12a. As *hes2.1* and *hes2.2* are tandem duplicates, we generated an approximately 30kb deletion spanning both genes. We refer to these double mutants as *hes2^-/-^*. *hes2^-/-^* neuromasts have a similar phenotype as *ccndx^-/-^* neuromasts with greater than 60% of hair cells showing a posterior polarity bias compared to 50% of sibling hair cells (Figure 7A-7C). Accordingly, Emx2^+^ hair cells are also increased in *hes2^-/-^* neuromasts (Figures 7E-7G), even though the total number of hair cells is unaffected (Figure 7D), demonstrating that *hes2* specifically affects hair cell polarity. Hence, *hes2* is necessary to establish proper hair cell polarity via negative regulation of Emx2 expression.

**Figure 7.**
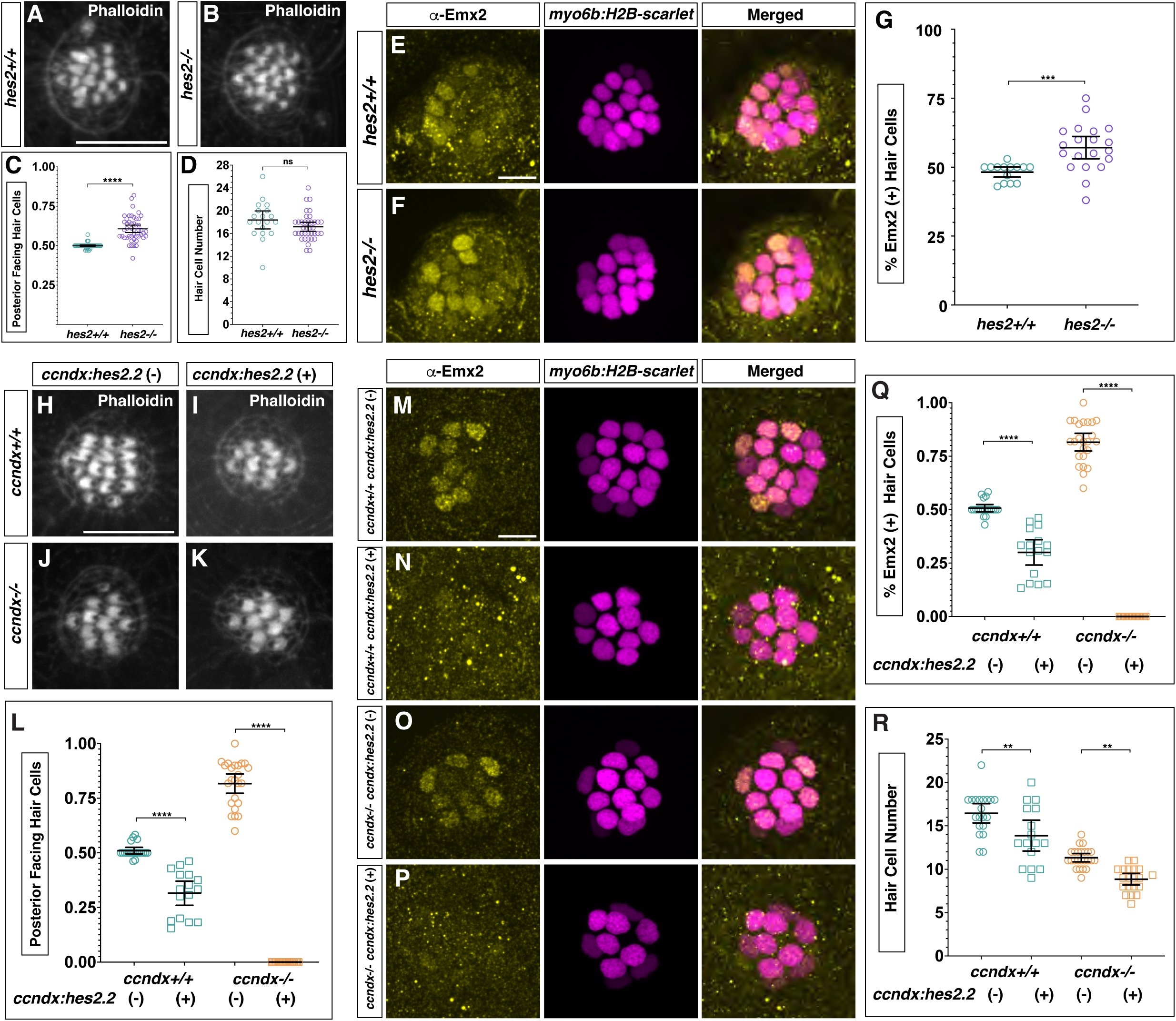
*hes2* is required for proper hair cell polarity through regulation of Emx2. (A-C) Phalloidin staining of 5 dpf neuromasts from sibling (A) and *hes2^-/-^* (B) with quantification of posterior facing hair cells per neuromast (C). Scale bar=10μm. (D) Quantification of hair cell number in 5 dpf neuromasts from sibling and *hes2^-/-^*. (E-G) Anti-Emx2 immunostaining (yellow) in *myo6b:H2B-mScarlet-I* (magenta) sibling (E) and *hes2^-/-^* 5dpf neuromasts (F). Scale bar=10μm. (G) Quantification of the percentage of Emx2^+^ hair cells per neuromast at 5 dpf. (H-K) Phalloidin staining of *ccndx:hes2.2,* 5 dpf neuromasts in *ccndx^+/+^* or *ccndx^-/-^*. Scale bar=10 μm. (L) Quantification of posterior facing hair cells per neuromast. (M-P) Anti-Emx2 immunostaining (yellow) in *ccndx^+/+^* or *ccndx^-/-^* neuromasts with or without *ccndx:hes2.2* expression. (Q) Quantification of Emx2^+^ hair cells per neuromast in *ccndx^+/+^* or *ccndx^-/-^* mutants with or without *ccndx:hes2.2* expression. (R) Quantification of hair cell number in in *ccndx^+/+^* or *ccndx^-/-^* mutants with or without *ccndx:hes2.2* expression.

To test the sufficiency of *hes2.2* in determining hair cell polarity, we generated a transgenic line in which the *ccndx* upstream region drives *hes2.2* expression (called *ccndx:hes2.2)*. We injected this transgene into a *ccndx^+/-^* outcross to test potential functional interactions. *ccndx:hes2.2* induces an anterior polarity bias in both wildtype and *ccndx^-/-^* (Figure 7H-7L). In *ccndx^-/-^* all hair cells are anteriorly polarized, whereas in siblings the polarity conversion is less severe. We speculate that either the lack of proliferation, or additional genes dysregulated in *ccndx^-/-^* neuromasts may affect *ccndx:hes2.2* activity. Matching the polarity phenotype, *ccndx:hes2.2* expression decreases Emx2 expression in both wildtype and *ccndx-/-* (Figure 7M-Q). Additionally, *ccndx:hes2.2* induces a small decrease in hair cell number (Figure 7R). We conclude that loss of *ccndx* leads to a reduction in *hes2* expression, causing an increase in Emx2, which results in posterior polarized hair cells.

Our combined results demonstrate that proliferation of amplifying stem cells and their progeny is independently regulated by different *cyclinD* genes. In addition, in the absence of proliferation, hair cells still fully regenerate, but hair cell polarity is affected due to the downregulation *hes2.2* and *emx2*.

## Discussion

Here, through transcriptional profiling and genetic approaches we show that in the zebrafish lateral line, proliferation of hair cell progenitors and amplifying stem cells is regulated by two different *cyclinD* family members, *ccndx* and *ccnd2a*, respectively. Organ homeostasis and regeneration depend on symmetric proliferation of stem- and progenitor cells to regenerate lost differentiated cells while maintaining both pools. A potential significance of individually regulating proliferation of different stem- and progenitor populations is that it enhances robustness of the regeneration process: Even if proliferation in one population is disrupted, the other still proliferates, leading to a certain level of short-term regeneration, which will, however, eventually come to a halt once the stem cells are depleted.

Our work raises the question if in other regenerating organs, such as the epithelia of the intestine, stomach, esophagus, the skin and the hematopoietic system, proliferation of stem cells and their progeny is also regulated by different *Cyclin D* genes. Some evidence exists that *Cyclin D* genes can have temporal or location-specific functions. For example, mouse developmental and adult neurogenesis depends on different *CyclinD* genes [48–52]. Region-specific expression of *CyclinD* genes also occurs along different segments of the mouse intestine [53]. However, to our knowledge it is unknown if stem cell and progenitor divisions are regulated by different *CyclinD* genes during the same neurogenesis stage or if distinct *CyclinD* genes regulate intestinal crypt or villus cells differentially, which would have intriguing therapeutic implications.

Cyclins can also have proliferation-independent functions. For example, CyclinD also affects developmental genes independently of association with CDK4/6, e.g. by recruiting histone acetyltransferases or deacetylases [41, 54–57]. In the mouse retina, CyclinD1 positively regulates Notch1 expression through recruitment of a histone acetyltransferase, thereby affecting Notch signaling [58]. This raises the question if *ccndx* and *ccnd2a* regulate distinct processes in addition to activating proliferation. Here we show that *ccnd2a* can substitute for *ccndx* and rescues both the *ccndx* mutant hair cell number and polarity defects. Therefore, *ccndx* does not possess any specific functions in hair cell production or polarity that *ccnd2a* does not possess.

Interestingly, *ccndx* is present only in anamniotes and has been lost during amniote evolution [35, 36]. *ccnd1 and ccnd2* genes and several *fgf* ligands have maintained synteny during evolution [59], which can be observed comparing human and zebrafish *ccnd1 and ccnd2* genes (Figure S7A-S7B). It appears that *ccndx* evolved from a duplication of this ancestral *ccnd*-*fgf* locus, and that amniotes lost *ccndx* and retained the other syntenic genes (Figure S7A-7C. Protein structure prediction shows structural conservation between zebrafish Ccndx, Ccnd1 and Ccnd2a (Figure S7D). We postulate that *ccndx* was lost in amniotes as other *cyclinD* genes took over its role. This hypothesis is supported by our finding that *ccnd2a* rescues *ccndx*^-/-^ mutant phenotypes.

The expression of cell fate specification genes is often associated with cell cycle regulatory mechanisms, as the cell cycle machinery can affect the epigenome, chromosome architecture and transcriptional programs required for cell fate. CyclinD genes and Cyclin-dependent kinases (CDKs) have been implicated in cell fate decisions in several cell types, and relevant to our study, CCND1/CDK phosphorylates and stabilizes Atoh1 in the mouse cerebellum [60]. Here we show that, even though *atoh1a* and *ccndx* possess highly similar expression dynamics and contrary to prior dogma, lateral line hair cell differentiation does not require proliferation, even though fewer hair cells regenerate. This finding mirrors what has been described in intestinal organoids where specification occurs prior to proliferation. However, proliferation rates control the abundance of differentiated intestinal cell types after specification. For example, different lineages divide at different rates leading to the generation of substantially more absorptive than secretory cells [61]. This raises the possibility that in the intestine proliferation is also regulated by different cell cycle genes in various cell lineages as we described for the lateral line system.

In contrast to hair cell differentiation, we show that the establishment of normal hair cell polarity does require progenitor proliferation in the lateral line. The prevailing model for establishment of neuromast hair cell polarity is that progenitor cell division generates two initially equivalent daughter cells, which then initiate Notch signaling between themselves. The posterior-facing hair cells express Emx2, and if overexpressed, Emx2 is sufficient to reverse the polarity of normally anterior-facing hair cells [28]. It is unknown what induces Emx2, but it is inhibited by Notch signaling [25, 30]. Stochastically, one daughter cell will have less Notch activity, become Emx2^+^ and posteriorly polarized [25, 28, 30, 46]. We found that *ccndx^-/-^* neuromasts possess more Emx2+ hair cells, and our scRNA-seq analysis revealed that *hes2.2* and *hey2* [62–65], among other genes, are downregulated in progenitor cells and young hair cells, respectively (Figure 6). These findings suggest that these genes could be involved in negatively regulating Emx2 expression, possibly downstream of Notch activation. However, in the mouse ear, Hey2 is not dependent on Notch signaling [66, 67]. Additionally, as in mouse cell lines or the frog retina [66–69], we determined that *hes2.2* is not a Notch target in neuromasts. *hes2.2* is therefore inhibiting Emx2 expression via a Notch-independent mechanism.

Our *hes2* mutation and overexpression experiments strongly suggest that *hes2* is acting downstream of *ccndx* to inhibit Emx2 expression. However, *ccndx* is expressed earlier and much more broadly than *hes2.2*, making direct regulation unlikely. On the other hand, *hes.2.2* is expressed before progenitor cells undergo cytokinesis but at the same time as mitotic genes such as *cdk1*. Thus, *hes2.2* appears linked to the cell cycle, but cell cycle events by themselves are insufficient to trigger *hes2.2,* as it is not induced in amplifying cells. As *hes2* is restricted to the hair cell lineage, it is likely regulated by hair cell progenitor specific genes that still need to be identified.

*hes2* is also expressed at low levels in the zebrafish and mouse ear [70–73], and it will be interesting to test if Hes2 plays an equally crucial role in setting up hair cell polarity in the inner ear of fishes and other species.

Notch signaling controls hair cell versus support cell specification via the process of lateral inhibition [15, 17, 18, 21]. During regeneration, downregulation of Notch signaling also leads to increased differentiation divisions [15, 17], which we demonstrate here, is entirely dependent on *ccndx*. The two roles of Notch in cell specification and proliferation are at least partially independent as hair cell differentiation does not depend on *ccndx* or proliferation. In addition, Notch signaling affects hair cell polarity by inhibiting Emx2 expression, even though the precise mechanism is not well understood [25, 30]. It is possible that cell proliferation and hair cell polarity are linked, either because loss of progenitor cell division causes defects in the asymmetric distribution of Notch pathway components into the two daughter cells or because *ccndx* directly regulates Notch pathway genes, as has been described for *Ccnd1* in the mouse retina [58].

In summary, our results demonstrate that proliferation of zebrafish lateral line stem and progenitor cells is independently regulated, with potentially important implications for our understanding of how, in other organs and species, the proliferation of quiescent and activated stem cells and their daughter cells can be manipulated when diseased. We also show that loss of hair cell progenitor proliferation leads to hair cell polarity defects via loss of *hes2*, which so far has not been implicated in hair cell polarity.

## Acknowledgments

We thank Dr. Nina Könecke for initial scRNA-seq analyses of the lateral line. We also thank members of the Stowers Institute Core facilities including Allison Scott, Kaitlyn Petentler and Anoja Perera from the Sequencing and Discovery Genomics Core, and Fang Liu and Jeff Haug from the Cytometry Core for expert technical assistance. We thank all members of the Stowers Institute Aquatics Facility for excellent zebrafish husbandry. We thank Drs. Robb Krumlauf and Aurelie Hintermann for comments on the manuscript and all members of the Piotrowski laboratory for helpful comments throughout this project. This research was supported by NIH (NIDCD) award 1R01DC015488-01A1 to T.P., funding from the Hearing Health Foundation to T.P., and institutional support from the Stowers Institute for Medical Research to T.P.

## Author contributions

M.E.L. and T.P. conceived the project, designed experiments and interpreted results. M.E.L., Y.T., D.M., J.P. and J.E.S. performed experiments and analyzed experimental data. S.C. performed computational analysis. M.E.L. and T.P. wrote the manuscript with input from all authors.

## Declaration of interests

The authors declare no competing interests.

## Supplemental Figures

**Supplemental Figure 1.**
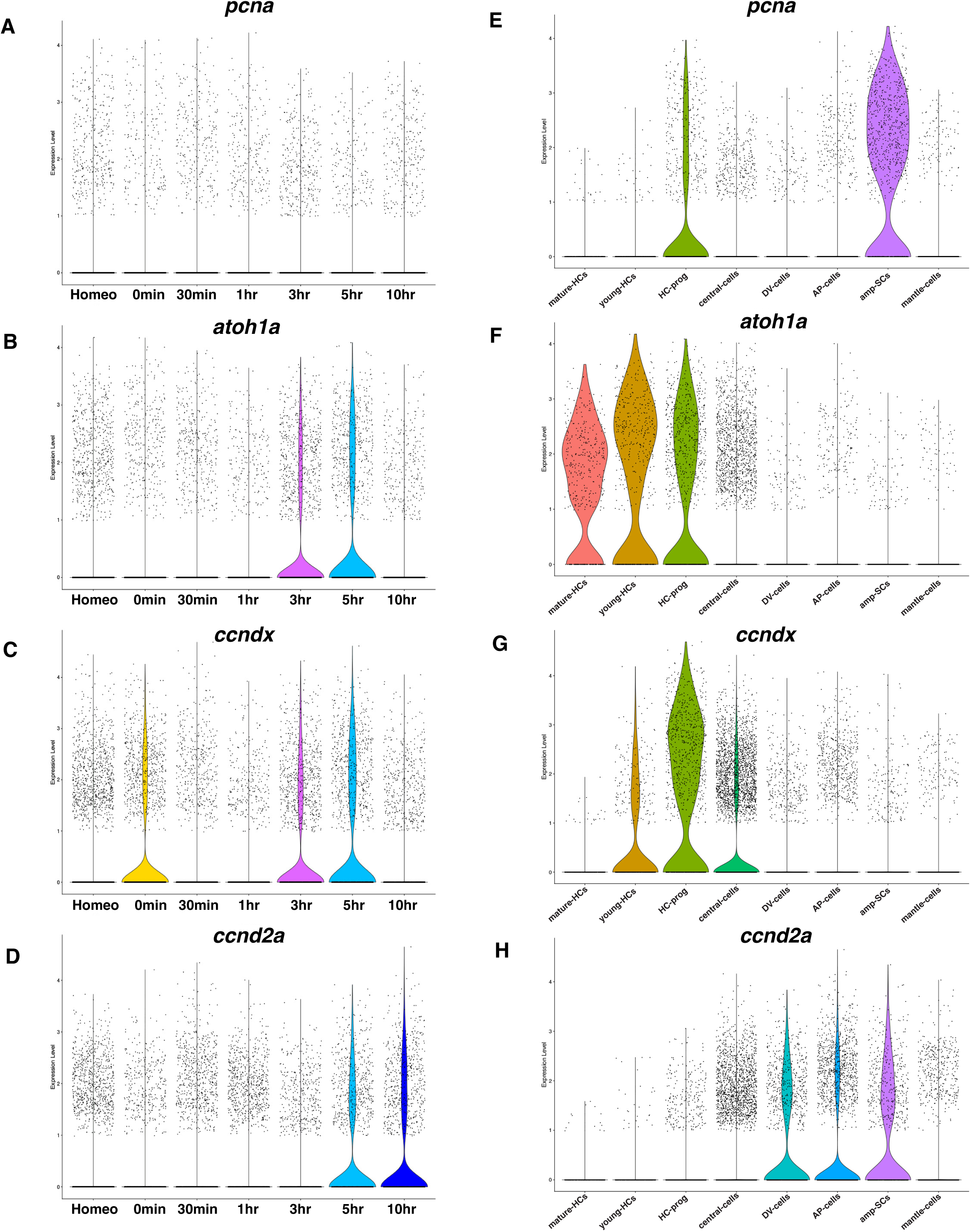
mRNA expression dynamics of genes in Figures 1G-K during regeneration from Baek et al., 2022 [12]. (A-D) Violin plots of *pcna*, *atoh1a*, *ccndx* and *ccnd2a* expression levels during the scRNA-seq regeneration time course (Baek et al., 2022) split by time point. (E-H) Violin plots of *pcna*, *atoh1a*, *ccndx* and *ccnd2a* expression levels during the scRNA-seq regeneration time course (Baek et al., 2022) split by cell type.

**Supplemental Figure 2.**
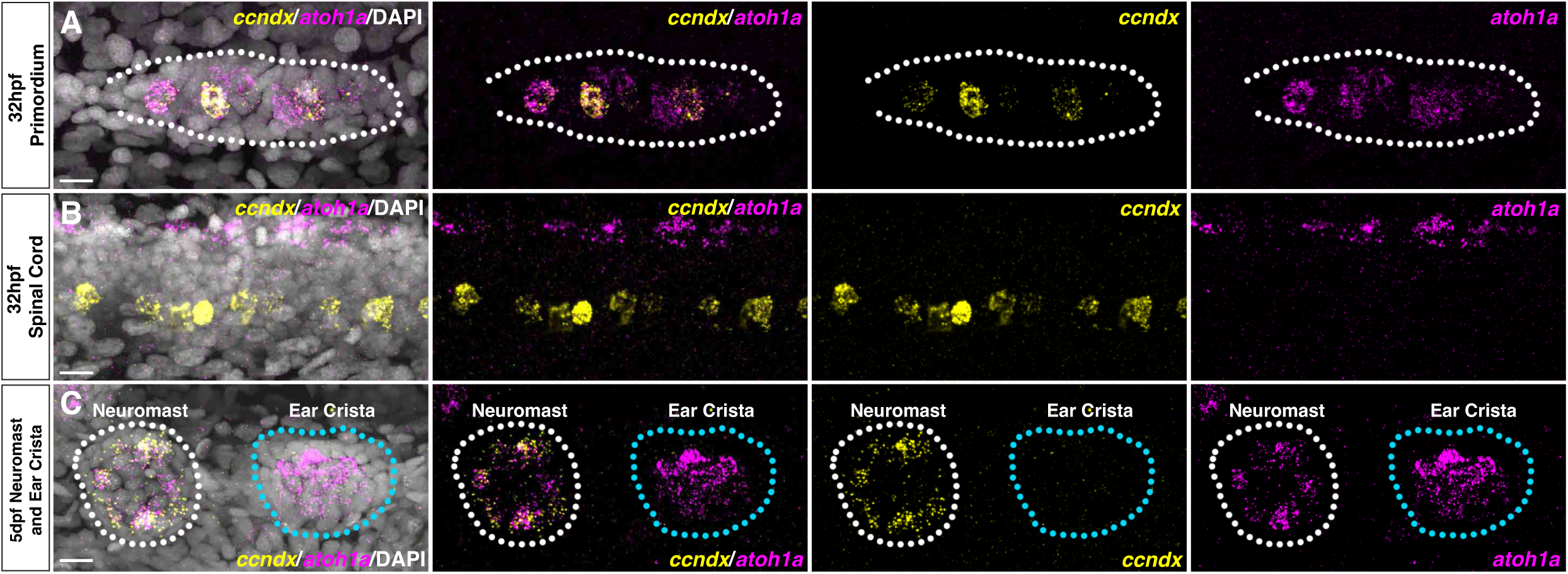
Additional regions of *ccndx* expression and comparison of *ccndx* with other *cyclinD genes* via synteny and protein structure prediction. (A) *ccndx* (yellow) and *atoh1a* (magenta) HCR in the 32hpf migrating lateral line primordium. Scale bar= 10 μm. (B) *ccndx* (yellow) and *atoh1a* (magenta) HCR in the 32hpf spinal cord. Scale bar= 10 μm. (C) *ccndx* (yellow) and *atoh1a* (magenta) HCR in a 5dpf anterior lateral line neuromast or medial crista. *ccndx* is not expressed in the crista. Scale bar= 10 μm.

**Supplemental Figure 3.**
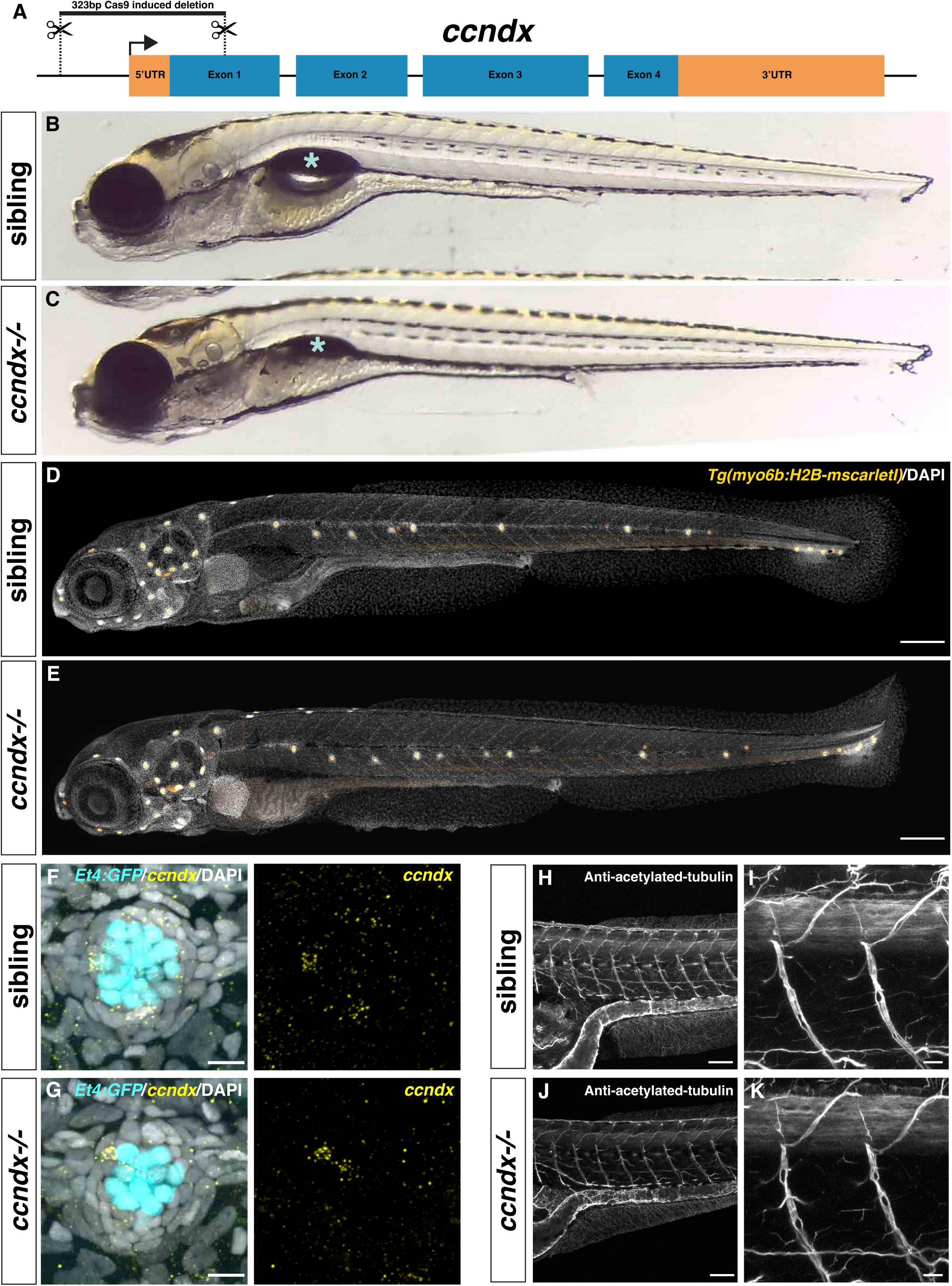
Generation and characterization of zebrafish *ccndx* mutants. (A) Diagram of *ccndx* showing the CRISPR-Cas9 induced 323bp deletion. (B and C) Widefield images of 5dpf sibling (B) and *ccndx^-/-^* (C). *ccndx^-/-^* lack an inflated swim bladder (asterisk), but otherwise appear morphologically normal. (D and E) Confocal images of 5dpf *myo6b:H2B-mScarlet-I* (orange) expressing sibling (D) and *ccndx^-/-^* (E) stained with DAPI (white) showing normal neuromast deposition in *ccndx^-/-^*. Scale bar= 200 μm. (F and G) *ccndx* HCR (yellow) in 5dpf *sqEt4:GFP* (cyan) expressing sibling (F) and *ccndx^-/-^* neuromasts (G). Scale bar= 10 μm. (H-K) Anti-acetylated tubulin immunostaining in 5dpf sibling (H-I) or *ccndx^-/-^* (J-K). Motor axons are present and appear normal in *ccndx^-/-^*. Scale bar= 100 μm in (H) and (J) and 10 μm in (I) and (K).

**Supplemental Figure 4.**
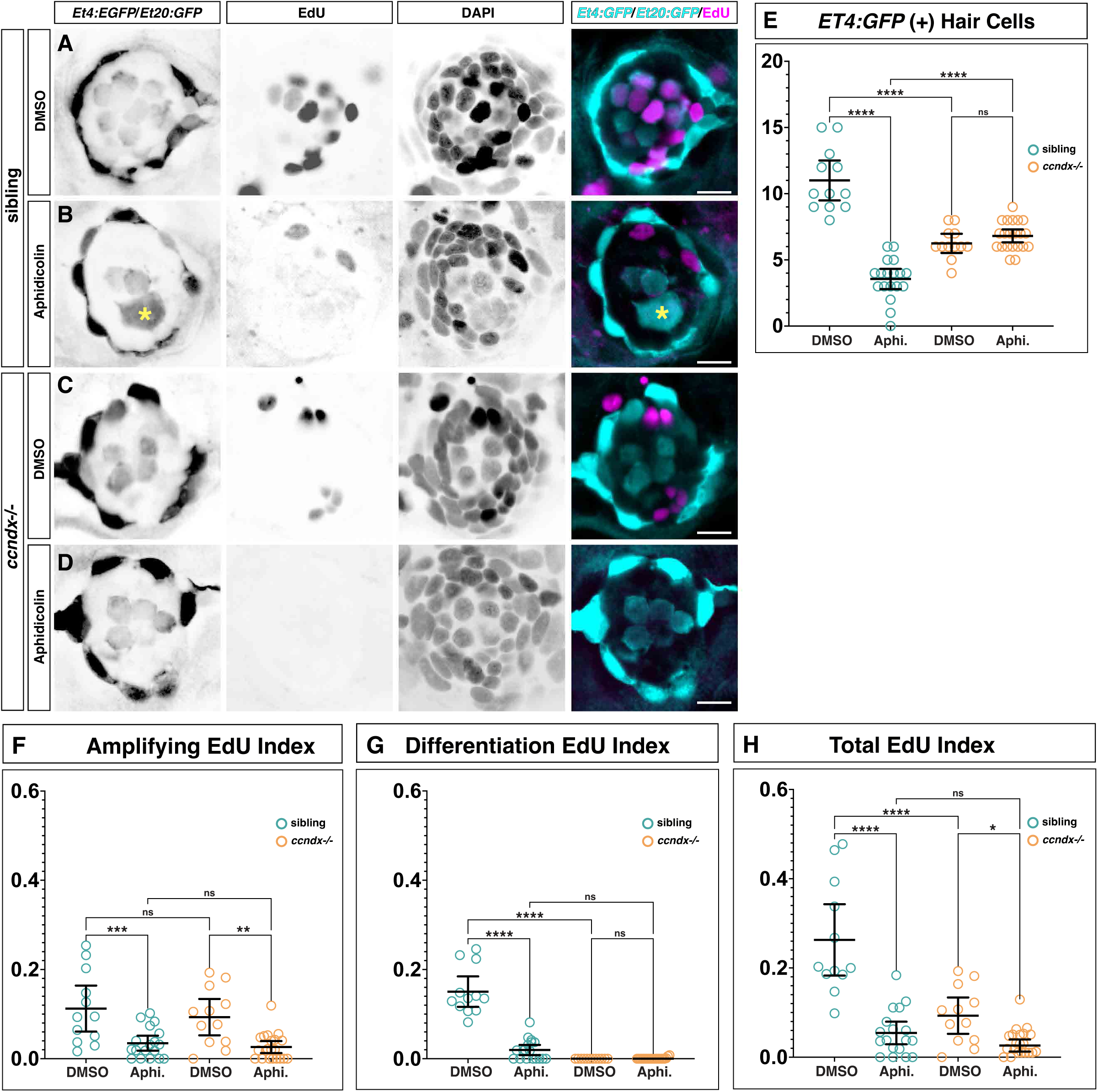
Aphidicolin-induced death of regenerating hair cells is rescued by loss of *ccndx*. (A and B) Representative images of EdU staining after 48 hrs of regeneration in DMSO- (A) and 100μM aphidicolin-treated sibling *sqEt4:EGFP/sqEt20:EGFP* larvae (B). An enlarged sq*Et4:EGFP*^+^ hair cell (asterisk) is seen after aphidicolin treatment. The two smaller hair cells in (B) likely survived neomycin treatment. Scale bar= 10 μm. (C and D) Representative images of EdU staining after 48 hrs of regeneration in DMSO- (C) or 100 μM aphidicolin-treated *ccndx^-/-^*/*Et4:EGFP/Et20:GFP* larvae (D). No enlarged *Et4:EGFP*^+^ hair cells are observed after aphidicolin treatment. Scale bar= 10 μm. (E) Quantification of *sqEt4:EGFP*^+^ hair cells in DMSO- or 100μM aphidicolin-treated sibling and *ccndx^-/-^* neuromasts after 48 hrs of regeneration. *ccndx^-/-^* form the same number of hair cells after DMSO or aphidicolin treatment. (F-H) EdU indexes from sibling and *ccndx^-/-^* neuromasts after 48hrs of regeneration in the presence of DMSO or 100 μM aphidicolin (Aphi). (F) Aphidicolin decreases the amplifying EdU index in both sibling and *ccndx^-/-^* neuromasts. (G) The differentiation EdU index in siblings is decreased after aphidicolin treatment likely due to hair cell death. In *ccndx^-/-^* neuromasts the differentiation index is the same after DMSO or aphidicolin treatment. (H) Total cell EdU indexes are decreased in both sibling and *ccndx^-/-^* neuromasts after aphidicolin treatment.

**Supplemental Figure 5.**
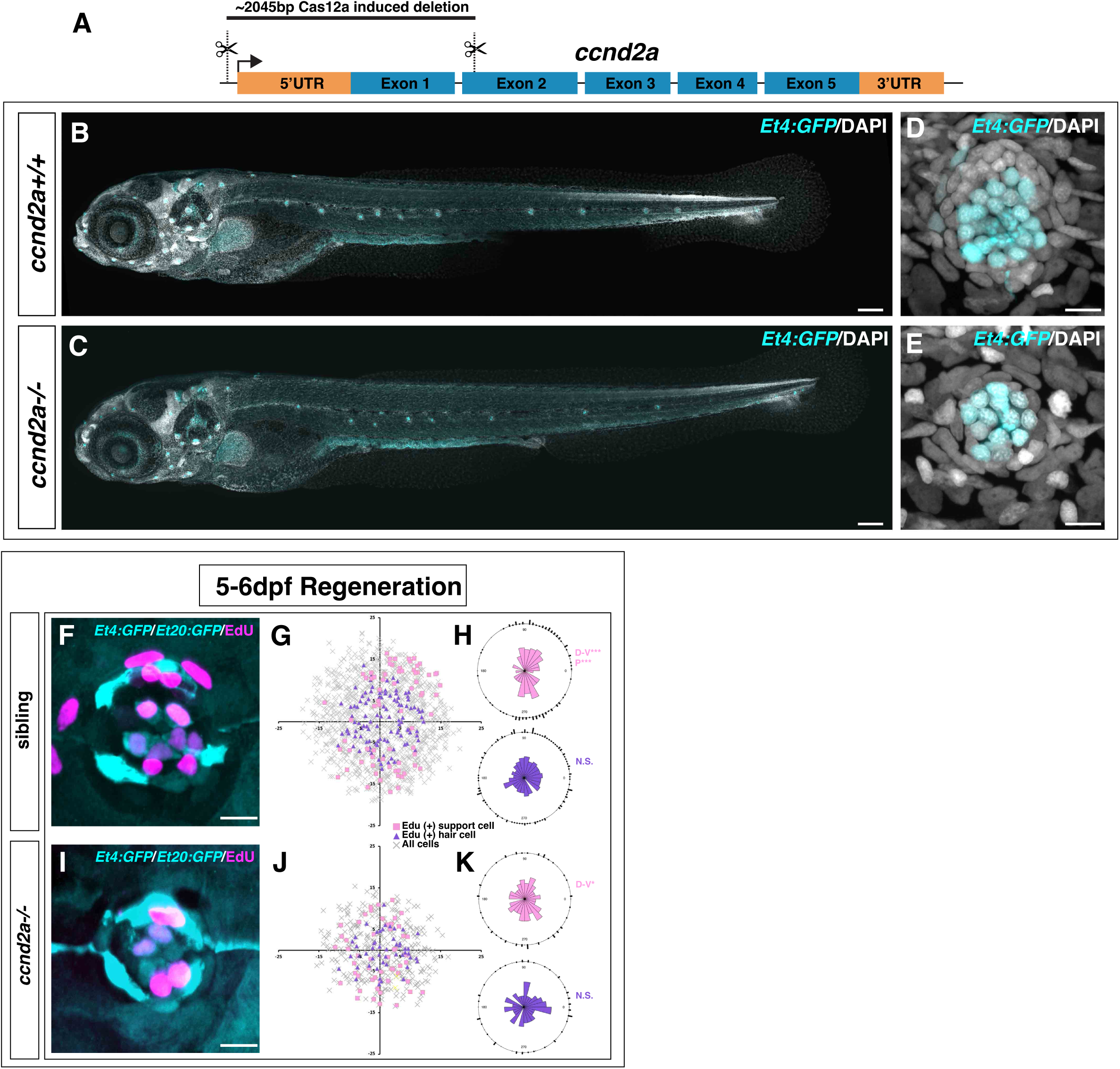
Generation of *ccnd2a* mutant zebrafish and characterization of regeneration phenotypes. (A) Diagram of the *ccnd2a* locus showing the CRISPR-Cas12a-induced deletion. (B-E) Confocal images of 5dpf sq*Et4:EGFP* (cyan) expressing wildtype (B and D) and *ccnd2a^-/-^* (C and E) zebrafish stained with DAPI (white) showing normal neuromast deposition in *ccnd2a^-/-^*. Scale bar= 200 μm in (B and C) and 10 μm in (D and E). (F-H) (F) Representative image of a sibling neuromast expressing *sqEt4:EGFP/sqEt20:GFP* stained with EdU (magenta) after regeneration from 5-6dpf. Scale bar= 10 μm. (G) Spatial analysis of multiple neuromasts of EdU^+^ cells. (H) Statistical analysis of positions of EdU^+^ support cells (pink) and hair cells (purple). (I-K) (I) Representative image of a *ccnd2a^-/-^* neuromast expressing *sqEt4:EGFP/sqEt20:EGFP* stained with EdU (magenta) after regeneration from 5-6dpf. Scale bar= 10 μm. (J) Spatial analysis of multiple neuromasts of EdU^+^ cells. (K) Statistical analysis of positions of EdU^+^ support cells (pink) and hair cells (purple).

**Supplemental Figure 6.**
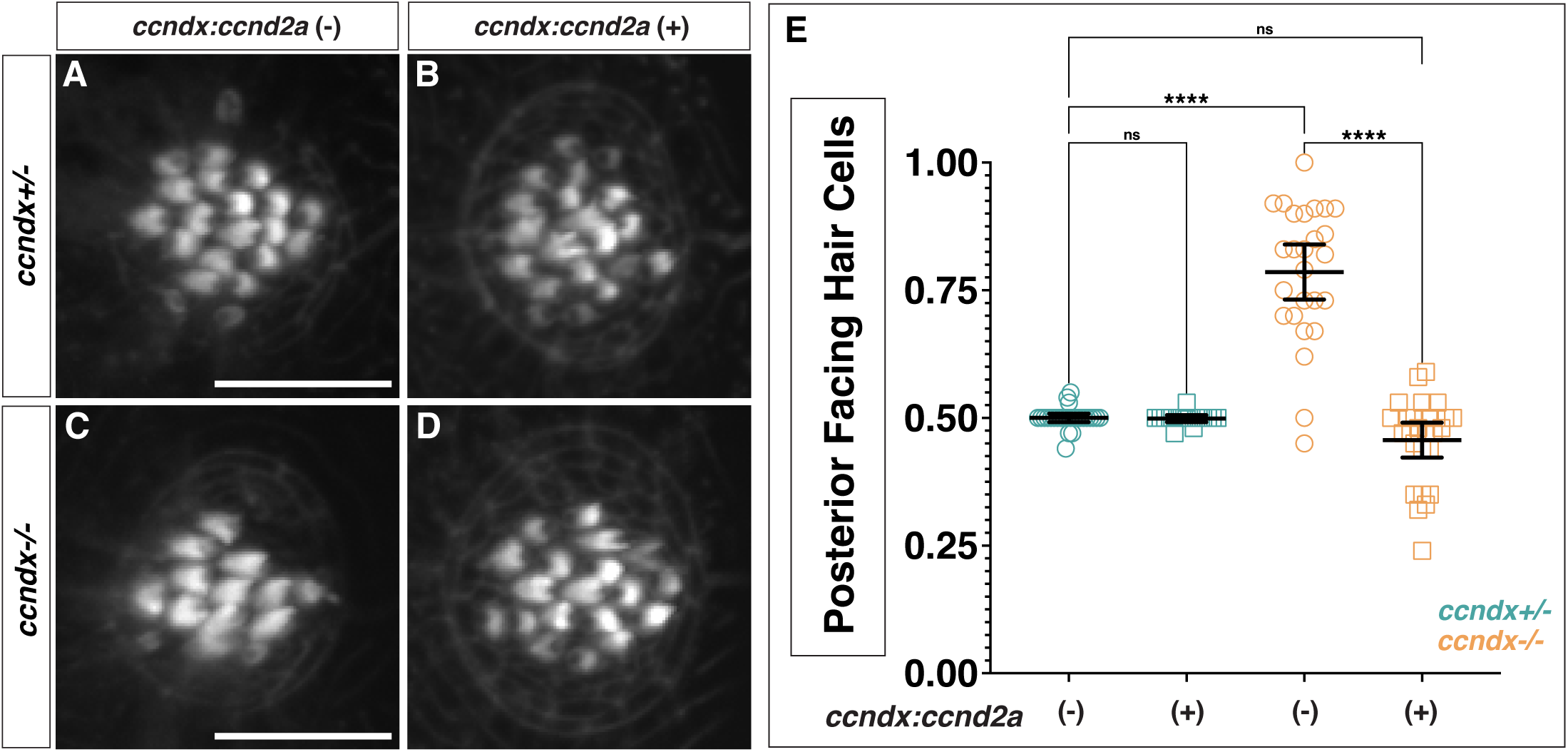
Expression of *ccnd2a* rescues the *ccndx^-/-^* hair cell polarity phenotype. (A-D) Representative images of phalloidin staining from 5 dpf neuromasts from *ccndx^+/-^* (A and B) or *ccndx^-/-^* (C and D) without or with *ccnd2a-P2A-mScarlet-I* driven by the *ccndx* promoter. Scale bar= 10 μm. (E) Quantification of posterior facing hair cells in *ccndx^+/-^* or *ccndx^-/-^* neuromasts without or with *ccnd2a-P2A-mScarlet-I*.

**Supplemental Figure 7.**
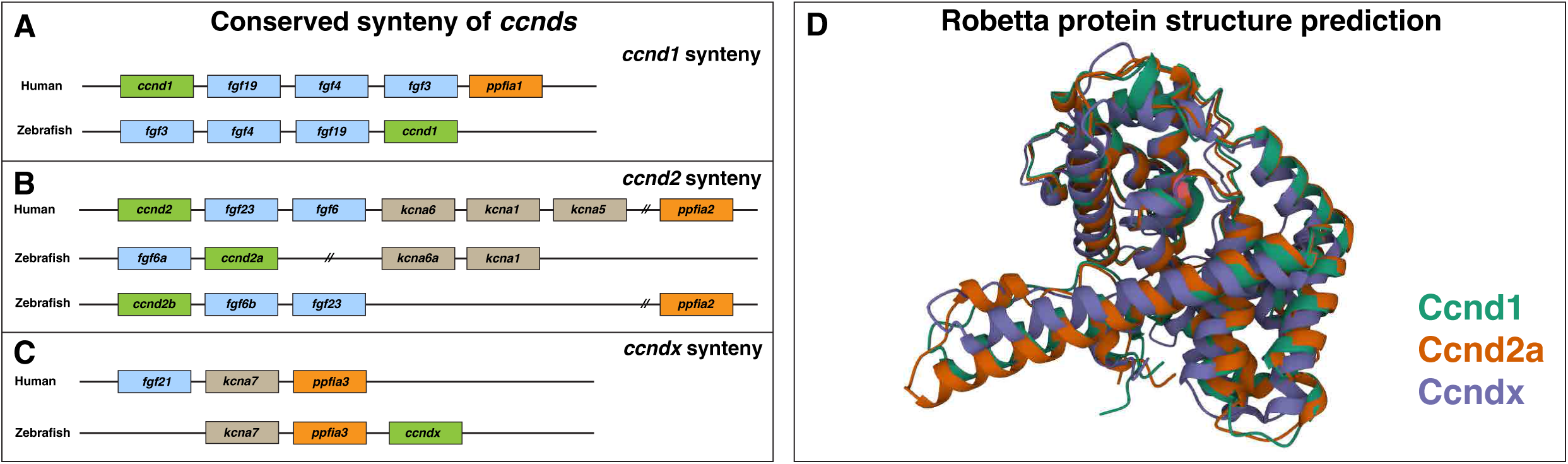
Evolution of *ccndx*. (A-C) Conserved synteny of Human and Zebrafish chromosome regions surrounding *ccnd1* (A), *ccnd2* (B) and *ccndx* (C). Zebrafish possess two copies of *ccnd2* and Humans lack *ccndx*. Adapted from Itoh and Ornitz, 2004 [59]. (D) Robetta protein structure prediction for zebrafish Ccnd1, Ccnd2a and Ccndx.

## Movie Legends

**Supplemental Movie 1. *ccndx^-/-^* neuromasts regenerate hair cells without progenitor cell division.**

Time-lapse recording of hair cell regeneration from 2-42 hrs post neomycin of sibling and *ccndx^-/-^* neuromasts expressing *myo6b:H2B-mScarlet*. In sibling neuromasts, *myo6b:H2B-mScarlet* starts to be expressed in progenitor cells as they divide. In *ccndx^-/-^* neuromasts *myo6b:H2B-mScarlet*^+^ hair cells form but never divide.

**Supplemental Movie 2. *ccndx^-/-^* hair cells are rescued from aphidicolin-induced death.** Time-lapse recording of hair cell regeneration from 5-44 hrs post neomycin of sibling and *ccndx^-/-^* neuromasts treated with aphidicolin immediately after neomycin. Larvae express both *ccndx:NLS-d2EGFP* and *myo6b:H2B-mScarlet*. In the sibling larva two hair cells survive and in the *ccndx^-/-^* larva three hair cells survive the neomycin treatment. In the sibling, new *myo6b:H2B-mScarlet-I*^+^ hair cells form but enlarge and eventually die. In the *ccndx^-/-^* neuromast new *myo6b:H2B-mScarlet-I*^+^ hair cells regenerate and survive through the end of the time-lapse.

**Supplemental Movie 3. Notch inhibition increases *ccndx:NLS-d2GFP* expression and generates more hair cells during regeneration.**

Time-lapse recording of hair cell regeneration from 2-41 hrs after neomycin in *ccndx:NLS-d2EGFP/myo6b:H2BmScarlet-I-* expressing neuromasts either left untreated or treated with the Notch inhibitor LY411575 immediately after hair cell death. In untreated zebrafish *ccndx:NLS-d2EGFP* becomes strong in progenitor cells, before they express *myo6b:H2BmScarlet-I* and then divide to generate two hair cells. *ccndx:NLS-d2EGFP* expression goes down as hair cells mature. In LY411575 treated fish *ccndx:NLS-d2EGFP* shows stronger expression after 10 hrs post neomycin and remains elevated in support cells throughout the time-lapse recording while decreasing in mature hair cells.

**Supplemental Movie 4. Notch inhibition during regeneration leads to the generation of more hair cells in *ccndx^-/-^* neuromasts without progenitor cell division.**

Time-lapse recording of hair cell regeneration from 2-42 hrs in *ccndx:NLS-d2EGFP/myo6b:H2BmScarlet-I* expressing neuromasts from sibling and *ccndx^-/-^* larvae treated with LY411575 immediately after hair cell death. The sibling neuromast shows increased hair cell generation from dividing progenitor cells. The *ccndx^-/-^* neuromast generates extra hair cells in the absence of progenitor cell division.

## Methods

### Zebrafish strains and husbandry

All zebrafish experiments were performed according to the guidelines established by the Stowers Institute for Medical Research IACUC review board.

### Zebrafish Mutants

*atoh1a^psi69^* [12] Primers Fw-GAGCAGAGCGAGTACCCACC and Rv-AGTTTCAGTTCCGACAGCTCG are used for genotyping followed by Sanger sequencing with the forward primer. *ccndx^psi76^* was generated via CRISPR-Cas9-induced mutagenesis using two single guide RNA (sgRNA) sequences AGCTGTTGTTGTGTCCCTCT**GGG** and CGGGACAGAGGGTCATTCAG**AGG** (PAM sites in bold) to generate a deletion from the upstream promoter region through exon one. sgRNA sequences were found using CRISPRscan [74]. sgRNA guides, without PAM sequences, and Cas9 tracrRNA were ordered from IDT. Guides were diluted to 100μM in Nuclease-free duplex buffer (IDT). To generate duplexes 1.5μL of each sgRNA and 3μL tracrRNA (100μM) were brought up to 50μl with dH_2_O, heated to 95^°^C for 5 minutes and left at room temperature for 20 minutes and stored at −20^°^C until use. For the injection mix 5μL of duplexed sgRNAs, 1μl Cas9 protein (1μg/uL, PNA BIO), 2μL 0.5% phenol red and 2μL dH2O were incubated at room temperature for 10 minutes, then kept on ice until injection. 1-3nL was injected into the cell of one-cell stage zebrafish embryos. Transient mutations were verified at 48hpf to prove that the sgRNAs worked and remaining embryos were raised to adulthood and screened for germline mutations. For genotyping, primers used were Fw-ATGGCTTTCAGATTGCAACA and Rv-CATTGCTCCAGGTTTTGGTT for the deletion mutation and Rv-CGCACCACAGAGACACAGAC with the same forward primer above for the wildtype PCR.

*ccnd2a^psi77^* was generated via CRISPR-Cas12a induced mutagenesis using single guide RNA sequences **TTTA**AAATGACAACACAAACTCAC and **TTTA**GCGGTGGTACCAACAAGAAA to generate a deletion from the upstream promoter region through part of exon two, which generated an approximately 2045b base pair deletion (PAM sites in bold). To find the guide sites, genomic regions were first screened via DeepCpf1 [75] to find scores above 10, then potential guides were BLASTed via the UCSC Genome browser to look for potential off target regions. Cas12a and guide RNA injections were performed based on previously published methods with modifications [76]. Cas12a guides without PAM sequences (IDT) were diluted to 24μM in dH_2_O. 100uM Lba-Cas12a protein was ordered from New England Biolabs. The injection mix consisted of 12μM guide mix, 20μM Lba-Cas12a, 0.3M KCl, 0.05% phenol red and brought up to 5μL with dH_2_O then kept for 10 minutes at 37°C. 1-3nL of this mix were injected into the cell of one-cell stage zebrafish embryos. Transient mutations were verified at 48 hpf and remaining embryos were raised to adulthood and screened for germline mutations. For genotyping, primers used were Fw-TGACACCAAGAGCATGGGTA and Rv-TGGACCCTTAAAAGCAGTAGGA for the deletion mutation and Rv-CGCATAAAGGGCTGAATGTC with the same forward primer above for the wildtype PCR.

*Df(Chr8:hes2.2,hes2.1)^psi89^* chromosomal deletion was generated via CRISPR-Cas12a-induced mutagenesis using single guide RNA sequences **TTTG**GGATGAGCTATGTATATTAT and **TTTG**GTTGAAGCGATCATCAATCAA (PAM sites in bold) located in the 3’ untranslated region of *hes2.2* and just upstream of *hes2.1*, respectively, which generated an approximately 30kb base pair deletion. sgRNA design and injections were performed as above for *ccnd2a*. To genotype the mutant allele primers Fw-CGTGGCTTGGTTAATTATTGCG and Rv-AAAATTATTGGCCCCTTTAAGC are used. Primers Fw-AAAGCGCTCATTCTCCCTCT and Rv-CATCATTCTGGAGCTCTGCG are used to genotype the wildtype allele.

### Zebrafish Transgenics

Previously published zebrafish transgenic lines used: *Tg(myo6b: hist2h2l-mScarletI)^psi66Tg^* [77], referred to as *myo6b:H2B-mScarlet-I* in the text. *Tg(Hsp70l:atoh1a)^x20TG^* [78], *Tg(she:H2A-mcherry)^psi57Tg^* [77], *Tg(tp1bglobin*:*EGFP)^um14Tg^* [79], *Et(krt:EGFP)^sqet4ET^* [80] and *Et(krt:EGFP)^sqet20ET^* [80].

### Zebrafish transgenic lines generated

*Tg(−4.5ccndx:hist2h2l-EGFP)*^psi78Tg^ (referred to as *ccndx:H2B-GFP* in the text) transgenic was generated as follows: a fragment 4.5kb upstream of *ccndx* was PCR’d from genomic DNA using primers aactcgagTGGAGGGTTTCTTGAACCTTT (forward) and aaggatccGTCCTGTTGCACGTGTGTCT (reverse) containing XhoI or BamHI sites respectively for cloning into the p5’-entry vector (p5E) from the tol2 kit [81]. This includes part of the 5’UTR of *ccndx*. PCR was performed with Phusion High-Fidelity DNA Polymerase (New England Biolabs) using zebrafish genomic DNA. Primers were ordered from IDT. The p5E-4.5*ccndx* plasmid was sequence verified via Oxford Nanopore sequencing by Plasmidsaurus. The p5E-4.5*ccndx* plasmid was used in a standard Gateway reaction using pMiddle entry H2B-EGFP, p3’entry-polyA [81] and pToneDest tol1 destination vector (Addgene plasmid #67691, [82]). Tol1 mRNA was generated using the T7 mMessage kit (Thermo Fisher Scientific) using linearized pToneTP as the template (Addgene plasmid #67692, [82]). To generate transgenics 1-3nl of an injection mix consisting of 12.5ng/μl DNA, 20ng/μl *tol1* mRNA, 0.3M KCl and 0.05% phenol red, was injected into the cell of a one-cell stage zebrafish embryos from a wildtype Tu incross. GFP^+^ larvae were screened at 48hpf and raised to adulthood then founders screened for germline transmission of GFP. For transgenic *Tg(−4.5ccndx:NLS-d2EGFP) ^psi81Tg^*, the sv40-NLS-d2GFP sequence from the pNSEN-d2 plasmid (Addgene plasmid #59763, [83]) was first cloned into the pMiddle-Entry plasmid. This was then used in a Gateway reaction using p5E-4.5ccndx, p3E-polyA and pToneDest. Injection and screening of founders was as described above.

For transgenic *Tg(−4.5ccndx:ccnd2a-P2A-mScarlet-I)^psi83Tg^* a *ccnd2a* expression construct gBlock was ordered (IDT) consisting of zebrafish *ccnd2a* followed by a P2A-mscarletI sequence. This gBlock was first cloned into the pCR4-topo cloning plasmid (Thermo Fisher Scientific) and sequenced verified. This plasmid was then digested with EcoRI and the subsequent *ccnd2a-P2A-mScarletI* fragment was cloned into the EcoRI site of the pME and sequenced verified. A gateway reaction was performed with p5E-*4.5ccndx*, pME-*ccnd2a-P2A-mScarletI*, p3E-*polyA* and the pToneDest Tol1 destination plasmid. 1-3nl of the injection mix, consisting of 20ng/μl DNA, 16ng/μl *tol1* mRNA, 0.3M KCl and 0.05% phenol red, was injected into the cell of a one-cell stage zebrafish embryos from an in cross of *ccndx^psi76^*/*sqEt4:EGFP*/*sqET20:EGFP*. mScarlet^+^ larvae were collected at 48hpf and raised to adulthood. Founders were initially genotyped for presence of the *ccndx^psi76^* allele, then screened for germline transmission of mScarlet-I fluorescence. Multiple founders were identified, and founders that had a moderate level of *ccnd2a-P2A-mScarletI* expression were characterized.

### Hair cell ablation and proliferation analysis

To ablate hair cells 5 days post fertilization (dpf) zebrafish were treated with 300mM neomycin (Sigma-Aldrich) for 30 minutes. Neomycin was then extensively washed out and fish were allowed to recover for varying time points depending on the experiment.

To label dividing cells EdU (Carbosynth) was added at 3.3 mM with 1% DMSO in 0.5X E2 fish water. Larvae were added to EdU media immediately after neomycin treatment for 24-48 hrs, depending on the experiment, then fixed in 4% paraformaldehyde overnight at 4°C. Staining was carried out as described previously [11]. To visualize GFP in GFP-expressing fish rabbit anti-GFP (1/400, Thermo Fisher Scientific) immunolabeling was performed. For fish expressing *myo6b:H2B-scarlet* no antibody staining was needed. Alexa Fluor 594-Azide or Alexa Fluor 647-Azide (Thermos Fisher Scientific) were used during the Click-it reaction. Cell counts, EdU indexes and spatial positioning was performed as described [15, 37].

### Hybridization chain reaction (HCR) in situ hybridization

HCR probes and amplifiers were purchased from Molecular Instruments. Zebrafish were fixed at least two days in 4% paraformaldehyde at 4^0^C, dehydrated in a graded series of methanol, then stored in 100% methanol at −20^0^C until staining. HCR was performed according to the manufacture’s protocol (Molecular Instruments) with minor changes as previously described [84]. Zebrafish were permeabilized with 100% acetone at −20°C for 20 minutes for 5-6dpf zebrafish or 10 minutes for 32hpf zebrafish. HCR Probes used were *ccndx-B1, ccndx-B4, atoh1a-B2, atoh1a-B5, pcna-B4, ccnd2a-B2, ccnd1-B4*, *d2EGFP-B2,* and *hes2.2-B5*. For *ccndx* and *atoh1a* double HCR only the *ccndx-B4* and *atoh1a-B5* combination worked. Amplifiers used were conjugated with either Alexa-647, Alexa-546 or Alexa-488.

### Phalloidin staining and Immunohistochemistry

Phalloidin staining was performed as described with either Alexa Fluor 647, Alexa Fluor 568 or Alexa Fluor 488 phalloidin (Thermo Fisher Scientific) [85]. For anti-acetylated tubulin staining zebrafish were fixed in 2% paraformaldehyde/1% Trichloroacetic acid for 10 minutes at room temperature, then washed in PBS/0.8% Tween-20 (PBSTw) followed by 5 minutes wash in dH_2_O, 7 minutes in acetone at −20°C, then washed again in PBSTw. Larvae were then blocked at least 1 hour in 10% normal goat serum (GEMINI BIO-PRODUCTS) in PBSTw. Anti-acetylated tubulin (Sigma T6793) was diluted to 1/1000 in blocking buffer and incubated overnight at 4°C. Larvae were then washed in PBSTw and incubated in secondary antibody, goat anti-mouse Alexa-546 (Thermo Fisher Scientific) 1/1000 in blocking buffer, overnight at 4°C. The samples were then extensively washed.

Immunohistochemistry against Emx2 (Trans Genic, K0609) was performed as described [29]. In some cases, this was done simultaneously with rabbit anti-GFP (Thermo Fisher Scientific) at 1/400 dilution overnight. Secondary antibody goat anti-mouse Alexa Fluor 546 (Thermo Fisher Scientific Scientific) was used. DAPI was used to visualize nuclei, followed by Alexa Fluor 647 phalloidin staining (Thermo Fisher Scientific).

### Protein analysis

Ccnd protein structure comparison was performed with Robetta protein structure prediction (https://robetta.bakerlab.org/) [86], using zebrafish Ccnd1, Ccnd2a and Ccndx amino acid sequences downloaded from Ensembl [44]. The protein structures were visualized and superimposed using Mol* 3D Viewer, https://www.rcsb.org/3d-view [87].

### Imaging

Images were taken on either a Zeiss LSM 780 confocal microscope or Nikon Ti2 Eclipse equipped with a CSU-W1 Yokogawa spinning disk. Post-processing of images was performed with ImageJ (Fiji) [88].

### Time-lapse imaging

Hair cells were first killed with neomycin as described above before mounting. Zebrafish anesthesia and mounting in low-melt agarose were performed as described [37]. Time-lapse imaging was performed with a Nikon Ti2 Eclipse with a Yokogawa CSU-W1 spinning disk head with a Hamamatsu Flash 4.0 sCMOS with a temperature-controlled stage set to 28.5°C. During each time-lapse experiment the L2 and L3 neuromasts from two fish were imaged. Z-stacks were taken every 5 minutes for up to 42 hours. For quantification of *myo6b:H2B-mScarlet-I* hair cells the time when fluorescence signal could first be observed was recorded.

### Drug treatments

The gamma-secretase inhibitor LY411575 (Selleckchem), dissolved in DMSO, was diluted to a final concentration of 50μM. Aphidicolin (Sigma-Aldrich), dissolved in DMSO, was used at 100μM for EdU analysis and at 60μM for time lapse experiments. DMSO only was used as a negative control for both experiments.

### Statistical analysis

All statistics and graphs were generated with GraphPad Prism 9 (version 9.4.1). For comparing two groups unpaired t-test was used. For experiments with multiple groups a one-way ANOVA with Tukey’s multiple comparisons test was used to test significance. A p-value smaller than 0.05 was considered significant.

### Single cell RNA-sequencing

#### Embryo dissociation and FACS

scRNA-seq was performed in duplicate using 5 dpf homeostatic *ccndx* sibling and *ccndx^-/-^* fish. Siblings were collected from a cross between *ccndx^psi76^*/*Tg(she:H2A-mcherry)^psi57Tg^* and *Tg(she:H2A-mcherry)^psi57Tg^* zebrafish and sorted for mCherry expression at 4dpf. Mutant larvae were collected from an incross of *ccndx^psi76^*/*Tg(she:H2A-mcherry)^psi57Tg^* and sorted for mCherry^+^ fish lacking an inflated swim bladder at 4 dpf. These larvae were then allowed to recover until 5 dpf, then processed for FACS. Approximately 400-500 larvae were collected for each replicate.

Embryo dissociation and FACS were performed similarly to previously published protocols with modifications [12]. 5 dpf larvae were anesthetized with MS-222, then split between two 70mm cell strainers (Falcon) and placed in 0.25% Trypsin-EDTA (Gibco) on ice. The larvae from each strainer were then added to a 5ml round bottom tube (Falcon) and the tissue was broken up by slowly pipetting up and down using a pipette man and a glass Pasteur pipette (VWR), keeping the tubes on ice. The total time between placing larvae in trypsin to finishing pipetting was approximately 8 minutes. The dissociated tissue was then filtered through a 70μm filter (BD Biosciences), pelleted for 5 min at 2000rpm at 4°C and the pellet was resuspended in 1xDPBS/0.04%BSA. The pelleting and resuspension in 1xDPBS/0.04%BSA was repeated two more times. After the final was, the pellet was resuspended in 0.5mL 1xDPBS/0.04%BSA. The cells were combined and filtered through another 70μm filter into a 5ml round bottom tube. Cells were then incubated in Draq5 (1/200 dilution) for 5 minutes on ice. Draq5^+^/mCherry^+^ cells were sorted by the Stowers Institute Cytometry Core Facility using a BD Influx Cell Sorter (BD Biosciences, San Jose, CA. USA), with cells collected in tubes containing 10μL 1XDPBS/0.04% BSA. Sorted cells were then spun down at 350rcf at 4°C for 2 min in a swinging bucket centrifuge with slow deceleration. Most of the supernatant was removed and the cells were resuspended in approximately remaining 100μL.

### 10X Chromium scRNAseq library construction

The cells were then given to the Sequencing and Discovery Genomics Department (Stowers Institute for Medical Research) for sequencing. Dissociated, sorted cells were assessed for concentration and viability using a Luna-FL cell counter (Logos Biosystems). Cells deemed to be at least 80% viable were loaded based on live cell concentration onto a Chromium X Series instrument (10x Genomics) and libraries prepared using the Chromium Next GEM Single Cell 3’ Reagent Kits v3.1 (10x Genomics) according to manufacturer’s directions. Resulting cDNA and short fragment libraries were checked for quality and quantity using a 2100 Bioanalyzer (Agilent Technologies) and Qubit Fluorometer (Thermo Fisher Scientific). With cells captured estimated at ∼6,800-9,600 cells per sample, libraries were pooled and sequenced to a depth necessary to achieve at least 30,000 mean reads per cell on an Illumina NextSeq 2000 instrument utilizing RTA and instrument software versions current at the time of processing with the following paired read lengths: 28*10*10*90bp.

### Single cell RNA-seq Data Analysis

Raw reads were processed using 10X Genomics software Cellranger v7.0. Reads were demultiplexed into Fastq file format using cellranger mkfastq with default parameters. Genome index was built under zebrafish danRer11, Ensembl version 102 and used cellranger count function for cell gene count matrix generation.

Feature count matrix generated by cellranger were then loaded into R package Seurat (v4.3). Here we have two replicates for both sibling and mutant populations. We first processed each sample individually for quality control. The cells with less than 500 UMI counts or more than 10% of mitochondrial expression were filtered out. Then top 2000 variable genes were selected to run PCA analysis for nearest neighbor graph and clustering (PC=40, res=0.8), as well as the UMAP visualization. Then all replicates and samples were merged based on Seurat’s CCA scRNA-seq integration method for all downstream analysis. Finally, Seurat’s FindAllMarkers function (ROC test) was used to generate marker genes for every cluster and annotated based on ZFIN database knowledge.

## Notes

### Competing Interest Statement

The authors have declared no competing interest.

